# tRNA expression and modification landscapes, and their dynamics during zebrafish embryo development

**DOI:** 10.1101/2024.01.30.575011

**Authors:** Tom Rappol, Maria Waldl, Anastasia Chugunova, Ivo L. Hofacker, Andrea Pauli, Elisa Vilardo

**Affiliations:** Centre for Anatomy & Cell Biology, Medical University of Vienna, 1090 Vienna, Austria; Department of Theoretical Chemistry, University of Vienna, 1090 Vienna, Austria; Vienna Doctoral School in Chemistry (DoSChem), University of Vienna, 1090 Vienna, Austria; Institute of Computer Science and Interdisciplinary Center for Bioinformatics, Leipzig University, D-04107 Leipzig, Germany; Research Institute of Molecular Pathology (IMP), Vienna BioCenter (VBC),1030 Vienna, Austria; Faculty of Computer Science, Research Group Bioinformatics and Computational Biology, University of Vienna, 1090 Vienna, Austria

## Abstract

tRNA genes exist in multiple copies in the genome of all organisms across the three domains of life. Besides the sequence differences across tRNA copies, extensive post-transcriptional modification adds a further layer to tRNA diversification. Whilst the crucial role of tRNAs as adapter molecules in protein translation is well established, whether all tRNA are actually expressed, and whether the differences across isodecoders play any regulatory role is only recently being uncovered. Here we built upon recent developments in the use of NGS-based methods for RNA modification detection and developed tRAM-seq, an experimental protocol and *in silico* analysis pipeline to investigate tRNA expression and modification. Using tRAM-seq we analysed the full ensemble of nucleo-cytoplasmic and mitochondrial tRNAs during embryonic development of the model vertebrate zebrafish. We show that the repertoire of tRNAs changes during development, with an apparent major switch in tRNA isodecoder expression and modification profile taking place around the start of gastrulation. Taken together, our findings suggest the existence of a general reprogramming of the expressed tRNA pool, possibly gearing the translational machinery for distinct stages of the delicate and crucial process of embryo development.

## INTRODUCTION

tRNAs are the essential adapter molecules mediating the translation of messenger RNAs in the decoding centre of the ribosome. Besides the crucial role in protein synthesis, new regulatory functions of tRNAs and derived fragments have been uncovered in recent years (reviewed in (1)). Yet relatively little is known about the actual tRNA expression landscape and eventual dynamics in different physiological and/or pathological conditions. tRNA genes are present in multiple copies, with repertoires in higher eukaryotes ranging from a few hundred (about 600 copies in the human genome) up to thousands (e.g. over 20000 in zebrafish) (2). These estimates are based on computational predictions, and to date it is not clear whether all putative tRNA genes are actually transcribed, and whether their expression levels are modulated. Noteworthy, the expression of ribosomal RNAs has been shown to be heavily regulated during development in vertebrates, with distinct sets of rRNA genes being expressed in the oocyte and early stages of embryonic development versus later embryonic stages and adult zebrafish (3,4). While similar findings were recently reported also for other non-coding RNAs (5–7), reliable data on tRNAs remain scarce and under-represented in transcriptomic studies.

The study of tRNA expression is complicated due to technical reasons and the biology of tRNAs themselves. Typically, the genome encodes multiple copies of tRNAs for each anticodon. These multiple copies can be identical in the sequence of the mature tRNA, or they can be isodecoders, i.e. have identical anticodon sequence but differ in the rest of the sequence by a few or a substantial number of nucleotides (2). When (small) RNAs are analysed by next generation sequencing (NGS), reads mapping to multiple sites in the genome, so called multimappers, are usually discarded from further analysis. Furthermore, reads originating from tRNAs that present a few mismatches can also lead to mapping artefacts, or be discarded when the alignment parameters are not suitably adjusted. As a consequence, standard NGS data analysis pipelines cannot be used for analysis of tRNAs.

One additional challenge in the study of tRNAs is the presence of abundant modifications. To date, about 170 RNA modifications have been described, most of which are found in tRNAs (8). With up to 20% of nucleotides modified at the sugar or base moiety, tRNAs are the most modified RNA molecules in the cell. Many of the described modifications, particularly those present at the Watson-Crick face of the nucleobase, interfere with reverse transcription and can cause early termination or sequence errors in the cDNA. These “errors” sum up in the diversity of the reads originating from the tRNA transcriptome, requiring an optimized analysis approach. In recent years, new NGS strategies actually harnessed those sequencing errors for detection of RNA modifications (reviewed in (9)). Furthermore, the availability of enzymes able to remove RNA methylations (which are the most abundant tRNA modifications (10)), new reverse transcriptases, and improved mapping strategies have enhanced our ability to investigate tRNA expression and modification (11–14). Still, to date only few studies have addressed the possible modulation of tRNA expression and modification, for example in different tissues (15) or in response to stress and nutrient availability (see review (16)). Remarkably, mutations in the majority of tRNA modification enzymes are associated to diseases, typically characterized by developmental disorders and phenotypes restricted to specific tissues (17). These observations suggest that the hypomodification of tRNAs or the consequent depletion of mature tRNAs in their correct, functional form may be particularly detrimental in specific developmental stages and/or cellular contexts.

Here we analysed the dynamics of tRNA expression and modifications during embryo development in vertebrates, using zebrafish (*Danio rerio*) as model organism due to its fast and well characterized embryonic development (18). We optimized previously published small RNA sequencing protocols and devised an improved *in silico* analysis pipeline to delineate the expression profile of nuclear-encoded as well as mitochondrial tRNAs during the early stages of zebrafish embryo development. Furthermore, we have profiled a wide array of different tRNA modifications. We show that tRNA expression levels vary during embryo development, and specific tRNA modifications are dynamic in a tRNA- and developmental stage-specific manner. Taken together, our results suggest the existence of a dynamic reprogramming of tRNA expression and modification during embryogenesis, possibly contributing to the fine-tuning of the protein synthesis machinery during development.

## MATERIALS AND METHODS

### Fish husbandry

Zebrafish (*Danio rerio*) were raised according to standard protocols (28 °C water temperature; 14/10-hour light/dark cycle). TLAB fish, generated by crossing zebrafish AB and the natural variant TL (Tupfel Longfin), were used for all experiments. All fish experiments were conducted according to Austrian and European guidelines for animal research and approved by the Amt der Wiener Landesregierung, Magistratsabteilung 58—Wasserrecht (animal protocols GZ 342445/2016/12 and MA 58-221180-2021-16).

### RNA and tRNA fraction isolation

Samples from individual stages were collected as follows: between 100-200 chorionated embryos were collected per time point at the indicated stages. Embryos were transferred into 1.5 ml tubes and as much water as possible was removed before embryos were homogenized in 500 µl of TRIzol (Invitrogen). To obtain eggs, females were anesthetized using 0.1% Tricaine and eggs were isolated via a standard protocol in zebrafish (squeezing) (19). Between 100-200 eggs were activated by adding fish water (5 mM NaCl, 0.17 mM KCl, 0.33 mM CaCl_2_, 0.33 mM MgSO_4_, 10−5% methylene blue) and incubated for 10 minutes before collection and homogenization in TRIzol. To obtain ovaries, females (one for each replicate) were dissected. Ovaries were harvested and homogenized in 500 µl TRIzol.

Total RNA was extracted using the standard TRIzol protocol. Total RNA was separated by denaturing polyacrylamide gel electrophoresis (PAGE), and the small RNA fraction (∼60-120 nt) containing tRNAs was eluted in elution buffer (1 mM EDTA, 0.5 M NH4OAc and 0.1% SDS), phenol–chloroform extracted and precipitated.

### Expression and purification of recombinant proteins

The expression vectors encoding truncated AlkB and AlkB D135S (13) pET30a-AlkB and pET30a-AlkB-D135S were a gift from Tao Pan (Addgene plasmid # 79051; http://n2t.net/addgene:79051; RRID:Addgene_79051, and Addgene plasmid # 79050; http://n2t.net/addgene:79050 ; RRID:Addgene_79050). pET30a-AlkB-D135S/L118V (20) was constructed by performing site directed mutagenesis on pET30a-AlkB-D135S using the QuikChange protocol (Agilent Technologies) and the oligos L118V_FW (GATTTCCAGCCAGATGCTTGT<u>G</u>TTATCAACCGCTACGCTCCT, mutation underlined) and L118V_RW (AGGAGCGTAGCGGTTGATAA<u>C</u>ACAAGCATCTGGCTGGAAATC, mutation underlined). The expression and purification protocol was described previously (21). Briefly, each expression vector was transformed in *E. coli* BL21(DE3) pLysS and grown shaking at 37 °C until an OD_600_ of 0.6-0.8. Expression was induced with 1 mM IPTG and 5 μM FeSO4 for 4 hours at 30 °C. Cleared bacterial lysates were run on HisTrap FF Crude 1 ml columns (Cytiva), followed by a 1 M NaCl wash (to remove nucleic acids bound to AlkB), elution, and concentration with Vivaspin 10,000 MWCO (Sartorius). The purity of the protein fractions was verified by SDS-PAGE followed by Coomassie staining (Supplementary Figure S1A). The concentrations were calculated by comparison with BSA standards on SDS-PAGE; the enzymes were diluted to 500 µM, snap-frozen, and stored in aliquots at -80 °C.

### AlkB demethylation activity assay

The activity of a mixture of purified AlkB, AlkB D135S and AlkB D135S/L118V was tested for demethylation activity on RNA oligonucleotides with a 5’ modification under conditions previously described (21), similar to the demethylation treatment to be used on zebrafish tRNA (Supplementary Figure S1B,C). In a reaction volume of 25 µl, 50-250 pmol of AlkB, AlkB D135S and AlkB D135S/L118V were mixed and incubated with 1 pmol of a synthetic RNA oligonucleotide carrying m^1^A or m^1^G at the 5’-end (m^1^AUGCACUUGGACGAACCAGAGUGUAGCUUAA, IBA Sciences; m^1^GGCGCAGCGGAAGCGUGCUGGGCCCA, kindly provided by R. Micura) previously ^32^P-labelled with T4 PNK (NEB), and 500 ng total RNA extracted from HAP1 cells, in demethylation buffer (50 mM HEPES KOH pH 8, 1 mM α-ketoglutaric acid, 2 mM sodium ascorbate, 75 μM (NH_4_)_2_Fe(SO_4_)_2_, 50 μg/ml BSA) (12) and 40 U Murine RNase Inhibitor (NEB). Aliquots were withdrawn after 3 and 30 minutes at 25 °C, stopped by adding guanidine hydrochloride (final 166 mM), and further processed for TLC as previously described (22). The integrity of the RNA was verified by denaturing PAGE and GelRed staining.

### Library preparation for Next Generation Sequencing

Demethylation: AlkB treatment was performed in reaction volumes of 50 μl in demethylation buffer (see previous section), 40 U Murine RNase Inhibitor (NEB), 0.75 μg Zebrafish tRNA, 500 pmol AlkB, 500 pmol AlkB D135S, and 500 pmol AlkB D135S/L118V for 30 minutes at 25 °C, followed by phenol:chloroform:isoamyl alcohol (25:24:1) extraction (PCI) and ethanol precipitation.

Bisulfite conversion: 500 ng of demethylated tRNA were treated with the EZ RNA Methylation Kit (Zymo Research) according to manufacturer’s instructions.

Deacylation and dephosphorylation: tRNA samples were deacylated by incubation at 37 °C for 30 minutes in 70 mM Tris-HCl pH 9. Deacylation was omitted for bisulfite converted tRNA. Each tRNA sample was dephosphorylated by incubation at 37 °C for 45 minutes in a volume of 100 µl containing 1X PNK Reaction buffer, 10 U of T4 Polynucleotide Kinase (NEB) and 40 U Murine RNase Inhibitor (NEB), followed by PCI extraction and ethanol precipitation.

3’-adapter ligation: the tRNA pellets were redissolved and a pre-adenylated adapter (rApp-NNNNNNCTGTAGGCACCATCAAT-ddC) (23) was ligated to the 3’-end by incubation in a reaction volume of 20 µl, containing 20 pmol 3’-adapter, 1X T4 RNA ligase buffer, 25% PEG-8000, 200 U T4 RNA ligase 2 (truncated KQ, NEB) and 40 U Murine RNase Inhibitor (NEB) for 4 hours at 25 °C. The ligation products were separated on a 10% denaturing polyacrylamide gel and gel pieces spanning the range of about 55 nt to 300 nt were excised, crushed, and eluted over night at room temperature. The eluted tRNA was PCI extracted and precipitated.

Reverse transcription (RT): the adapter-ligated tRNA pellets were redissolved and annealed in a volume of 12 μl to 2.5 pmol RT primer (p-NNNAGATCGGAAGAGCGTCGTGTAGGGAAAGAGTGTAGATCTCGGTGGTCG C-Spc18-CACTCA-Spc18-TTCAGACGTGTGCTCTTCCGATCTATTGATGGTGCCTACAG) (23) by 2 minutes of incubation at 85 °C followed by 5 minutes of incubation at 25 °C. The reverse transcription was carried out in a reaction volume of 20 μl containing the annealed tRNA-RT primer, 500 nM TGIRT-III (InGex, or kindly provided by A. Lambowitz), 1X Protoscript II buffer (50 mM Tris-HCl pH 8.3, 75 mM KCl, 3 mM MgCl_2_) 40 U Murine RNase Inhibitor (NEB), 5 mM DTT and pre-incubated for 10 minutes at 42 °C. Then, dNTPs were added to a final concentration of 1.25 mM, followed by incubation at 42 °C for 16 hours. The remaining RNA in the RT sample was hydrolysed by adding 1 μl 5 M NaOH and incubation of 95 °C for 3 minutes. The cDNA was loaded on a 10% denaturing polyacrylamide gel and gel pieces were excised above the unextended RT primer till 400 nt, crushed, and the cDNA was eluted two times for 1 hour at 70 °C, 1500 rpm in 325 µl 1X TE pH 8. The two elutions were pooled and isopropanol precipitated.

cDNA circularization and amplification: the recovered cDNA was circularized by incubation in a reaction volume of 10 μl in the presence of 50 U CircLigase™ ssDNA Ligase (Epicentre), 1 M betaine, 1X CircLigase buffer, 50 μM ATP and 2.5 mM MnCl_2_ for 3 hours at 60 °C, followed by 10 minutes at 80 °C to deactivate the enzyme. Amplification of 4 µl circularized cDNA was performed in a reaction volume of 48 μl in the presence of 500 nM PCR forward primer (AATGATACGGCGACCACCGAGATCTACA*C, where * is a phosphorothioate bond) and indexed NEBNext® Multiplex Oligoes for Illumina (NEB), 0.48 U KAPA HiFi Polymerase (KAPA Biosystems), 1X KAPA HiFi GC Buffer and 30 μM dNTP Mix with an initial denaturation at 95°C for 3 minutes, followed by 13 to 15 cycles of 98 °C for 20 seconds, 62 °C for 30 seconds and 72 °C for 30 seconds. Libraries were purified with DNA Clean&Concentrator-5 kit (Zymo) or GeneJET Gel Extraction and DNA Cleanup Micro Kit (Thermo Fisher Scientific), quantified using Qubit dsDNA HS Assay (Invitrogen) and sequenced on a NextSeq500 instrument (Illumina) at the Core Facilities of the Medical University of Vienna, a member of VLSI.

### tRNA NGS data analysis

Details on the NGS data analysis can be found in Supplementary File 1. The results of the whole analysis are available in Supplementary Data 1. Here in the following is a short summary of the strategy and steps performed.

Preprocessing: The NGS data sets were preprocessed by trimming adapters, extraction of UMIs, trimming of untemplated nucleotides at the 5’ end, size filtering and quality filtering.

Reference genome: The initial tRNA reference genome was created by collecting mitochondrial tRNA sequences from mitotRNAdb and genomic (nuclear) sequences from a tRNAscan-SE run provided in GtrnaDB, including low confidence and pseudogene predictions. All DM samples were mapped to this initial reference genome (segemehl) and any genes with a coverage of at least 500 rpm in at least one sample were included in the reference genome. The reference genome was further refined manually. Canonical positions were manually assigned to the RFAM tRNA alignment (RF0005) and then transferred to the reference genome by aligning the tRNA references to the RFAM alignment.

Mapping: Mapping was performed with segemehl with optimized parameters for short reads. Bisulfite treated samples were mapped with the bisulfite version of segemehl (-F1) and RNA specific post-processing was applied.

Clustering: References were clustered based on (i) sequence similarity and (ii) by multimapper counts obtained from mapping. In the first step any references that differ in at most 3 positions were merged. In a second step, any clusters that consisted of more than 50% reads that can also be found in another larger cluster were merged into this larger cluster. The mapping results were merged for all references within one cluster.

Abundance: The abundance of each cluster was computed based on the DM samples by mapping on the manually refined reference set, random assignment of multi-mapping reads to one of the best matching references, counting of the mapped reads per reference, summing up the abundance of all references in a given cluster and normalizing the abundance count by the total count of mapped reads.

Misincorporation rates: Possible modification sites and possible changes of modification level were identified by computing position-wise misincorporation rates. For each reference position, the total number of reads that cover the given position were counted. Furthermore, we counted how many of those reads mismatch at the given position. The overall misincorporation rate per position in the clusters was computed by first adding up the mismatch counts and total coverage count for equivalent positions in the tRNA references within the given cluster, and then dividing the mismatch count by the total count. Equivalent positions between tRNA references within a cluster were defined by alignment against the tRNA RFAM alignment (RF0005).

RT stop fraction: RT stops were analysed by computing the fraction of reads that start immediately after a given position. The RT stop fraction for each position was calculated dividing the number of reads that start at one nucleotide downstream of the position of interest by the total number of reads that cover this downstream position in the reference tRNA. The overall RT stop fraction of a position within a cluster was computed by summing all read-starts that map to one position downstream of the cluster position of interest and dividing by the number of reads that cover this downstream position in the cluster.

m^5^C fractions: Putative m^5^C modification sites and m^5^C modification dynamics were detected based on the BS samples as C-retention rate. For single tRNA references, the C-retention rate was only computed for C positions and the count of reads that contain a C in the mapped position was divided by the count of mapped reads that contain a C or a T in the given position. To derive C-retention rates per cluster and per position, the count of mapped Cs was obtained as the number of reads with a C that is mapped to a reference C; the count of mapped Ts was obtained as the number of reads with a T that is mapped to either a reference C or T. The C-retention rate was computed as the count of Cs divided by the sum of the counted Cs and Ts. For interpretation details see also supplementary text in Supplementary File 1.

### Gene expression analysis

mRNA sequencing was performed using QuantSeq 3’ mRNA-Seq Library Prep Kit (FWD) for Illumina (Lexogen). 500 ng total RNA per sample were used. Single-indexed QuantSeq libraries were QC-checked on a Bioanalyzer 2100 (Agilent) using a High Sensitivity DNA Kit for correct insert size and quantified using Qubit dsDNA HS Assay (Invitrogen). Pooled libraries were sequenced on a NextSeq500 instrument (Illumina) in 1×75bp single-end sequencing mode. On average 8 million reads were generated per sample. Reads in fastq format were generated using the Illumina bcl2fastq command line tool (v2.19.1.403). Reads were trimmed and filtered using cutadapt (version 2.8) to trim polyA tails, remove reads with N’s and trim bases with a quality of less than 30 from the 3’ ends of the reads. On average, 7 million reads were left after this procedure. The composition of the reads after each trimming step was assessed using FASTQC (v0.11.9). Reads in fastq format left after the last trimming step were aligned to the *Danio rerio* (zebrafish) reference genome version GRCz11 with Ensembl 104 annotations using STAR aligner (version 2.6.1a) in 2-pass mode. Reads per gene were counted by STAR. Raw read counts were normalized and variance stabilizing transformation (vst) was done with DESeq2 (version 1.22.2). The mean vst-counts for known tRNA modifying enzymes (genes) per timepoint were plotted in a heat-map. Genes of interest were clustered and sorted corresponding to a UPGMA tree based on their similarity in expression levels as implemented in scipy as hierarchical linkage clustering method (scipy.cluster.hierarchy.linkage with method=’average’ and metric=’euclidean’).

### Codon frequency analysis

The codon frequency analysis was based on the same QuantSeq data as the gene expression analysis. Raw read counts were normalized by total number of mapped reads and converted to reads per million (RPM). Coding sequences were obtained from Ensembl (https://ftp.ensembl.org/pub/release-107/fasta/danio_rerio/cds/) (24). A representative gene isoform was selected based on primary isoform annotations in APPRIS 2023_01.v48 database (https://appris.bioinfo.cnio.es, zebrafish assembly version GRCz11, gene dataset Ensembl 104) (25,26). For each protein coding gene, the occurrences of each codon was counted and multiplied by the genes expression level (normalized read count). To obtain the codon frequency, those per codon counts were summed up over all genes and then divided by the total number of codons.

### Northern blotting

Approximately 2 µg of zebrafish total RNA were separated on a 15% denaturing polyacrylamide gel and imaged after GelRed staining. After pilot experiments, the amount of total RNA for different samples were adjusted to have comparable amount of tRNA loaded per lane. The RNA was transferred to a Hybond-N+ Nylon transfer membrane (Cytiva) with a Trans-Blot SD blotting apparatus (BioRad) followed by crosslinking at 0.12 J/cm^2^ in a CX-2000 UV crosslinker (Analytik Jena). The membrane was then rinsed with 6X SSC and prehybridized in 6X SSC, 10X Denhardt’s solution and 0.5% SDS rotating at 40 or 42 °C. The following antisense oligonucleotides were ^32^P-labelled with T4 PNK (NEB) and used as probes, hybridized at the indicated temperature overnight: anti-Asp-GUC-1 GCGGGGATACTTACC (40 °C), anti-Asp-GUC-2 GCGGAGATACTGTCC (40 °C), anti-Lys-CUUACTGAGCTAGCCGGGC (40 °C), anti-iMet-CAUTCTGGGTTATGGGCCCA (42 °C), anti-mtPhe TGGTGCATGCGGAGCTTA (40 °C), anti-mtVal TGGTCAGGACGATCCGAT (40 °C). The next day the membrane was washed once for 15 minutes with 6X SSC and 0.1% SDS and two times for 15 minutes in 4X SSC and 0.1% SDS. The membrane was exposed to a Storage Phosphor Screen and signals acquired on a Typhoon scanner (Cytiva). Membranes were reused after stripping of the probe by incubation 3 times for 15 minutes at 75 °C in stripping solution (40 mM Tris-Cl pH 7.5, 0.1X SSC and 1% SDS). Bands were quantified by densitometry with Image Lab 6.1 (Bio-Rad). The bands’ intensity was normalized by the intensity of the tRNA band in the polyacrylamide gel stained with GelRed prior to blotting.

### Primer-extension assay

Primer extension experiments to detect modifications were performed as previously described (27) with minor adjustments. The method is based on the synthesis of cDNA using a radioactively labelled primer that anneals a few nucleotides downstream from the modified RNA residue of interest. The primer is extended using an RT sensitive to modification, which results in chain terminations in correspondence of modifications (as opposed to the highly processive, modification-tolerant TGIRT which rather introduces a mismatch). The primer was annealed in a volume of 2.5 µl containing 1 pmol primer (PheGAA-A14 TTCAGTCTAATGCTCTCCCA; GluCUC-A58 TGGTTCCCTGACCGG ), 0.5-1.0 µg total RNA, 50 mM Tris-HCl pH 8.3, 60 mM NaCl and 10 mM DTT at 75 °C for 10 minutes followed by slowly cooling down to 23 °C. Then 2.5 µl containing dNTPs (10 mM), 10X Primer extension buffer and 2 U AMV Reverse Transcriptase (Promega) were added to the annealing mixture and incubated for 1 hour at 42 °C. The reaction was stopped by adding 5 µl of 2X RNA loading buffer. 5 µl of reaction were separated on a 15% denaturing polyacrylamide gel. The gel was exposed to a Storage Phosphor Screen at -80 °C. The RT stop fraction for the modification assayed was determined by dividing the band intensity of the stop observed immediately at the 3’ of the modification by the sum of the downstream bands intensity (consisting of stops due to other modifications or read-through to the RNA end. All the bands were quantified with Image Lab 6.1 (Bio-Rad).

### Mitochondrial DNA genotyping

After RNA extraction, DNA of the zebrafish samples ovary, 1k-cells, and bud was recovered from the TRIzol interphase/organic phase using back extraction buffer (4 M guanidine thiocyanate; 50 mM sodium citrate; 1 M Tris, pH 8.0) as previously described (28). Mitochondrial DNA was amplified with primers mtThr_F (GGAATAGCATTCCGCCCAG)/mtThr_R (GGTGGTCTCTCACTTGATATGGTG), mtAsp-Ser_F(CCGCCAAACGAGAAGTTCTG)/mtAsp-Ser_R(CCGGAAGGACTGTTCATACG), and mtArg_F(GCGGATTTGATCCACTAGGG)/mtArg_R(CACAGGCAGAAAAGGCTAGTAG). The amplicons were purified using the GeneJET PCR Purification Kit (Thermo Fisher Scientific) according to manufacturer’s protocol and verified by Sanger sequencing using the following primers: Thr GCAGACATGCTTGTACTAAC, Asp-Ser GTAGAATGATTACACGGCTG, Arg CCGCCTACCATTTTCATTACG.

## RESULTS

### tRAM-seq: an optimized library preparation and analysis protocol for tRNA profiling

To analyse tRNA expression and modification during embryo development, we isolated RNA from activated zebrafish eggs, as well as from embryos at the 4-cell stage (approximately one hour post-fertilisation, 1 hpf), 1000-cell stage (1k-cell, 3 hpf), 5 hpf, bud stage (10 hpf), and 24 hpf (Supplementary Figure S2); furthermore, we included samples of ovaries from adult female zebrafish.

For the preparation of sequencing libraries and their bioinformatic analysis, we built upon recent advancements in the use of NGS for similar purposes and devised an optimized <u>tR</u>NA <u>a</u>bundance and <u>m</u>odification <u>seq</u>uencing (tRAM-seq) protocol (Figure 1). The presence of RT-interfering modifications, the most abundant being methylations at the Watson-Crick face of the nucleobase (10), can result in under-representation in NGS data of the most heavily modified tRNAs. Similar to others (12,13), we used demethylation to minimize such bias and improve mapping, coverage, and quantification of the tRNAs. We used three variants of the demethylase of bacterial origin AlkB that were shown to have robust demethylase activity on *N*^1^-methyladenosine (m^1^A), *N*^3^-methylcytidine (m^3^C), *N*^1^-methylguanosine (m^1^G), and *N*^2^,*N*^2^-dimethylguanosine (m^2,2^G) (13,20,29). Thus, for each sample we isolated the tRNA fraction and split it in (i) a fraction not subject to demethylation (mock sample) used for modification detection (see following sections), (ii) a fraction treated with the mixture of the three demethylases used for tRNA quantification (DM sample), and (iii) a fraction subject to demethylation and bisulfite conversion for detection of m^5^C modification (BS sample) (Figure 1A).

**Figure 1.**
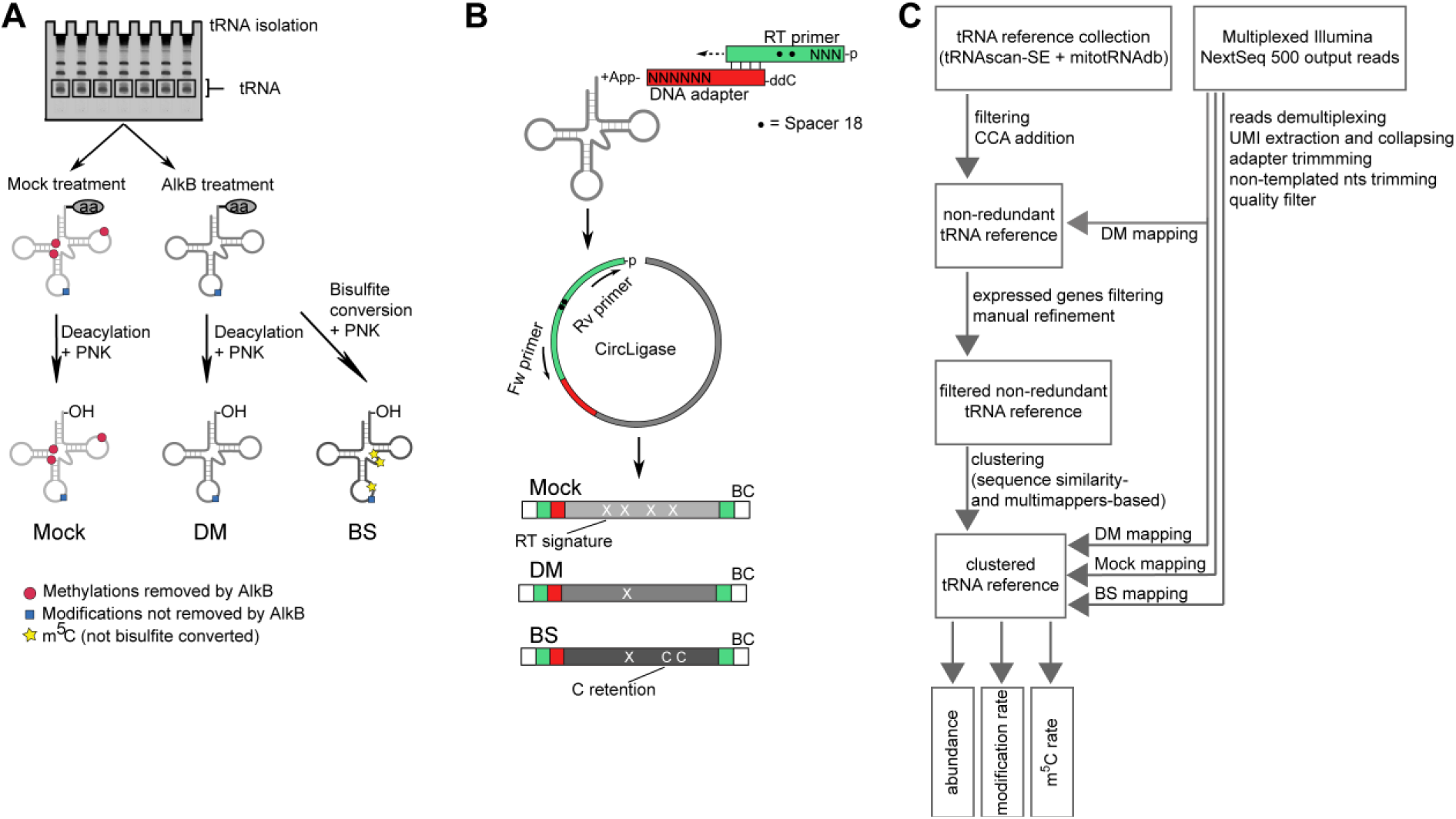
Library construction and computational workflow of tRAM-seq. (**A**) The tRNA fraction is isolated from total RNA and subsequently subject to treatments for demethylation (AlkB treatment), bisulfite conversion, and end repair (deacylation and T4 PNK treatment). **B**) The tRNA fraction is ligated to the pre-adenylated adapter (red) and reverse transcribed starting from the RT primer (green). Subsequently the cDNA is circularized and amplified from the RT primer sequence. The RT primer contains two internal spacers (•) to prevent circular amplification. Both the adapter and RT primer contain random nucleotides (N) at the 5’-end to reduce ligation bias and remove PCR duplicates during computational analysis. (**C**) Computational pipeline of tRAM-seq data analysis.

For library preparation, we adapted a protocol previously developed for small RNA sequencing (23), and further optimized it for tRNA sequencing (Figure 1B). We ligated the end-repaired tRNA samples to a pre-adenylated adapter, which bears six randomized nucleotides at the 5’-end in order to minimize sequence-preference bias in the ligation step. It was previously shown that different RTs are differentially affected by the modifications and structure of the template RNA, in terms of both processivity and fidelity (30). The thermostable group II intron reverse transcriptase TGIRT (31), with reaction conditions improved in a more recent report (11), was shown to be among the most processive enzymes: it is able to proceed past structure and modifications usually acting as roadblocks for reverse transcription, leaving a characteristic misincorporation signature at modified sites (30). We reverse transcribed the adapter-ligated RNA with TGIRT, and then circularized the synthesized cDNA with the 5’-end of the RT primer, which also includes three randomized nucleotides to minimize ligation bias. We amplified the obtained circular cDNA and sequenced the library by Illumina sequencing. The randomized nucleotides included in the library construction (6 in the ligated adapter and 3 in the RT primer) were used as unique molecular identifiers (UMIs) to remove PCR duplication products.

The presence of multiple, identical or nearly identical copies per tRNA gene requires optimized mapping strategies for the analysis of tRNA sequencing data. We and others have previously tackled the issue by using clustered references to map the generated reads (11,32–34), where multiple identical or nearly identical tRNA genes are collapsed to one representative reference. However, previous work did not address the case of the model organism zebrafish, which poses a major challenge of its own, with currently 8676 predicted high confidence tRNA genes, and additional thousands of lower-scoring predictions (2). Noteworthy, additional sequence variation can also be introduced by the presence of SNPs within laboratory zebrafish populations. After preliminary analysis, we found that our NGS data included reads from tRNA genes that are not part of the high-scoring GtRNAdb set. Therefore, we opted to include all predicted ∼20000 genes (including low-confidence predictions and pseudo genes) and performed mapping of the DM libraries on this initial set of tRNA candidate genes (Figure 1C). All tRNA genes represented by at least 500 reads per million (rpm) were included in our reference for further analysis, and after further filtering (see methods and Supplementary File 1 for details) the references were collapsed to 68 distinct clusters defined by sequence similarity and extent of multimapping reads (see methods and Supplementary Table 1 for details). The 68 clusters also included the 22 mitochondria-encoded tRNAs, which are all present in single copy in the organellar genome, corrected for SNPs that we identified in the sequencing data and that we confirmed by Sanger sequencing of amplified mitochondrial DNA (see supplementary text in Supplementary File 1 and Supplementary Figure S3).

All libraries were mapped to the refined, manually-curated clustered reference, and coverage and error rate were calculated for every position in the tRNA clusters’ sequences. In Figure 2 we show example coverage plots of the tRNA cluster Gln-CTG/TTG of a zebrafish ovary sample. The cluster includes reference tRNA genes differing at a few positions, namely canonical 34, 44, and 69, as indicated in the reference composition. Any other deviation from the expected sequences are flagged and highlighted. In the mock sample, the expected modifications m^1^G9 and m^1^A58 are clearly detected as a high error rate (Figure 2A). In the DM sample, nearly all signs of methylation at those positions were abolished, and only low, background-level errors are visible throughout the tRNA sequence (Figure 2B). The extent of full-length tRNA reads increased slightly in the DM versus the mock library, suggesting that, although our library preparation was efficient in the mock sample, the demethylation still improved the coverage toward the 5’-end of the tRNA cluster. The coverage plot for the BS library shows detection of m^5^C methylation at position 49 and 50 (Figure 2C), as expected for these tRNAs in vertebrates (8). Only a low level background of C retention is visible throughout the tRNA cluster sequence, indicating efficient bisulfite conversion. Similar results were obtained for all 68 tRNA clusters (Supplementary Files 2,3,4), thus showing that our tRAM-seq protocol enables us to assess tRNA abundance and modification in an organism with a very complex tRNA repertoire like zebrafish.

**Figure 2.**
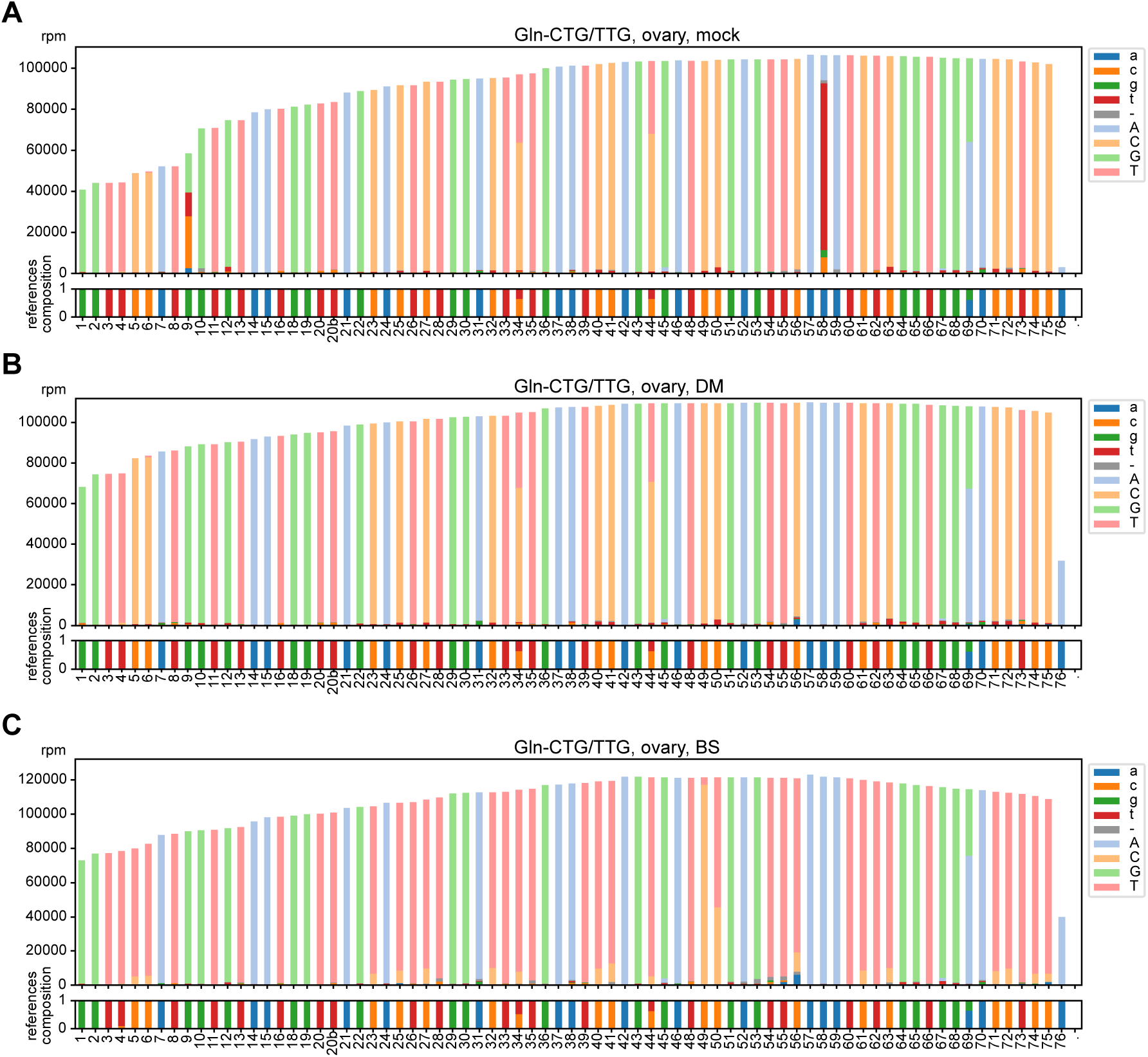
Coverage plots with modification signature. Example coverage plots with reference nucleotide identity of the cluster Gln-CTG/TTG for RNA isolated from zebrafish ovary and processed as (**A**) mock sample, (**B**) DM sample or (**C**) BS sample. Light colours indicate correctly mapped bases, dark colours highlight misincorporated bases, and grey highlights deletions. In (C), light orange colour represents C retention, indicative of m^5^C methylation. Numbers on the x-axis indicate canonical nucleotide position, and the amount of reads per million are shown on the y-axis.

### tRNA isodecoder identity and abundance are dynamic during embryo development

To profile the levels of tRNAs during early embryonic development, we used our tRAM-seq coverage data obtained from the DM samples. As shown in Figure 3A, we observed a wide range of expression levels for the different tRNA clusters, where tRNA-Asp-GTC was the most abundant, representing up to 10% of the total tRNA reads. Within the tRNA transcriptome, we observed diverse dynamics in tRNA levels across the samples analysed, with the expression of some tRNA clusters increasing or decreasing during embryogenesis, while other tRNA clusters appeared stable (Figure 3A,B). Noteworthy, we observed that the amount of total tRNAs as fraction of total embryo RNA (mostly consisting of rRNA) was highest in ovaries but lowest in activated eggs, and increased over the time course analysed until 24 hpf, (Supplementary Figure S4A). The fraction of mitochondrial tRNAs reads appeared to be highest in activated eggs and early embryo stages, and decreased over time (Figure 3A,B and Supplementary Figure S4B). Consequently, the apparent decrease in mitochondrial tRNAs was reasonably due to the prevalent increase of nucleo-cytoplasmic tRNAs rather than an actual decrease of the mitochondria-encoded ones. Normalizing the individual mitochondrial tRNAs to the mitochondrial pool instead of the total tRNA repertoire, we observed little changes in the relative abundance of the individual organellar tRNAs (Supplementary Figure S4C).

**Figure 3.**
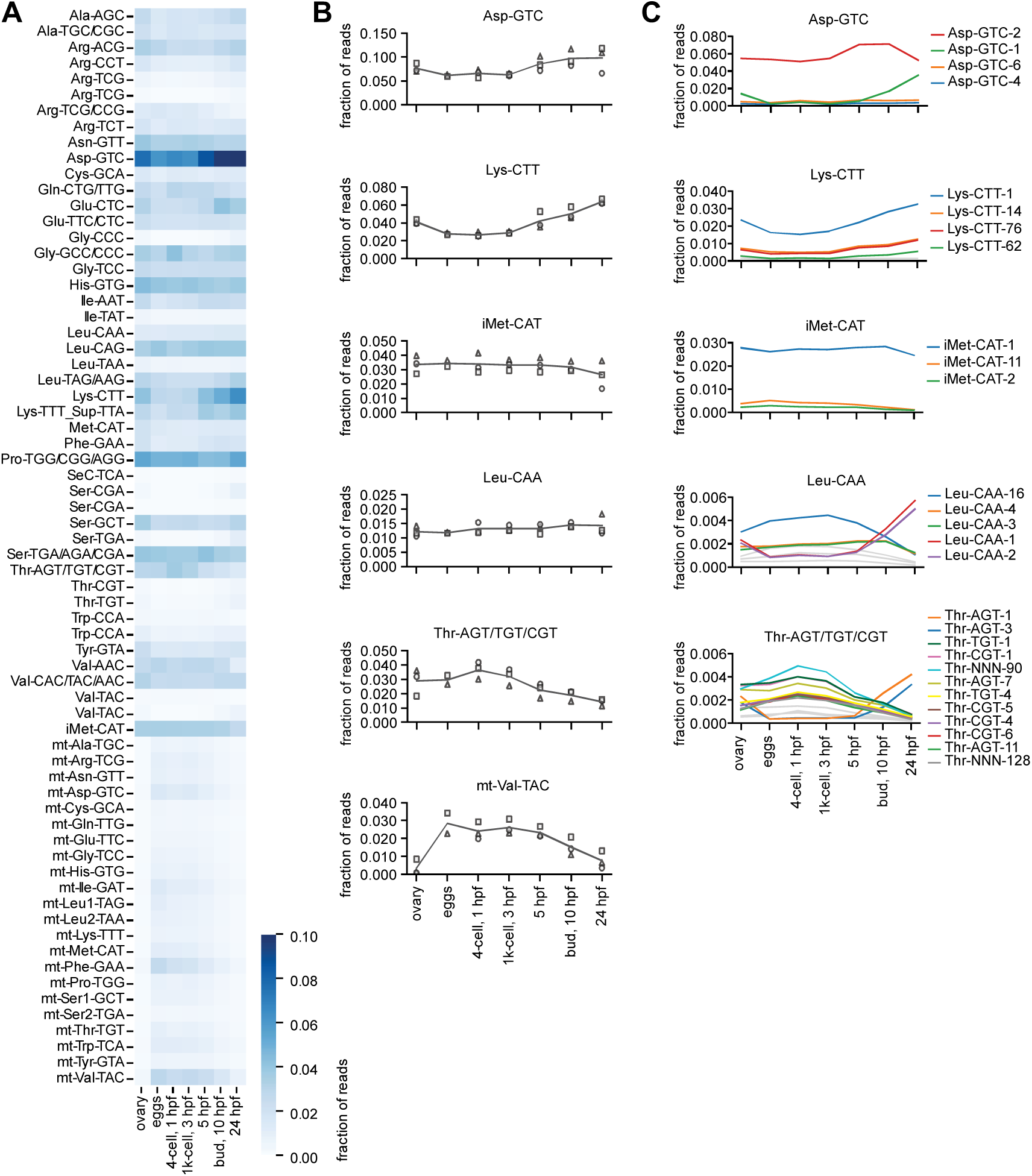
tRNA abundance during zebrafish embryo development. (**A**) Heat-map representing the normalized abundance of clustered nucleo-cytoplasmic and mitochondrial tRNAs (y-axis) in the analysed DM samples spanning zebrafish embryo development, plus ovaries (x-axis). The total number of reads mapped to the tRNA cluster were normalized by the total read count, and plotted per sample and time point. The blue colour scale indicates the fraction of reads mapped to the individual tRNA clusters. Data are means of three biological replicates with the exception of the activated eggs (n=2). (**B**) Abundance dynamics of selected nucleo-cytoplasmic and mitochondrial tRNA clusters. The line represents the mean of the three biological replicates with the exception of the activated eggs (n=2). Replicates are indicated by square, circle and triangle. (**C**) Plots showing abundance dynamics within tRNA clusters. Individual tRNA references are labelled and plotted in coloured lines if representing a minimum fraction of mapped reads of 0.002, the rest being plotted as grey lines. The line represents the mean of the biological replicates like in (B).

By deconvoluting the tRNA clusters into the individual tRNA isodecoders, we observed that in most instances a few tRNA isodecoders were the main representatives of each tRNA cluster. Remarkably, for a subset of tRNA clusters we observed an identity change between early and later time points, with a substantial switch in expressed tRNA isodecoders taking place from 5 hpf onward, as exemplified in the case of the clusters tRNA-Asp-GTC, Leu-CAA, and Thr-AGT/TGT/CGT (Figure 3C). Similar results were observed for other tRNA clusters, although not for all clusters (Supplementary File 5).

To validate our tRNA abundance results obtained by tRAM-seq, we chose northern blotting as orthogonal method. Northern blot relies on probe hydridization and is not affected by the same potential biases in ligation, RT, and PCR amplification that can affect NGS library preparation. In addition, we have designed our probes to target regions within the tRNA sequences which are devoid of modifications, aiming to prevent any bias due to differential modification extent. As shown in Figure 4, northern blot experiments on selected, representative tRNA clusters recapitulated closely the dynamics observed in the tRAM-seq data (Figure 3). Remarkably, using isodecoder-specific probes and stringent hybridization/wash conditions, we apparently distinguished the isodecoders tRNA^Asp-GTC-1^ and tRNA^Asp-GTC-2^, and confirmed their expression dynamics in the time course analysed.

**Figure 4.**
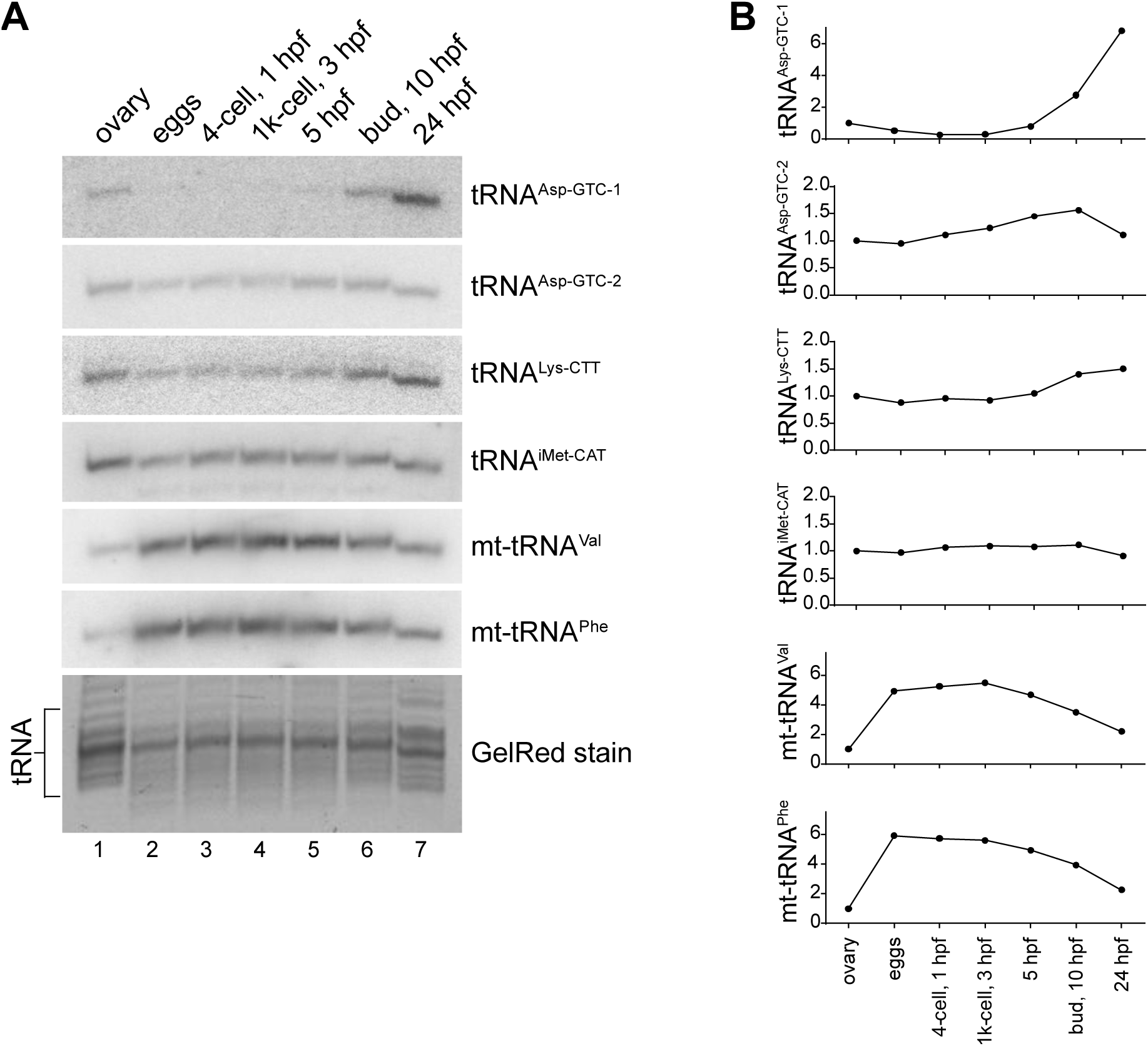
Quantification of tRNA abundance by northern blot. (**A**) The expression of selected tRNAs was analysed by northern blotting (upper six panels). The denaturing polyacrylamide gel was stained by GelRed before blotting and imaged to be used as loading control (bottom panel). (**B**) Quantification of tRNA abundance from northern blots (A) calibrated according to the GelRed-stained tRNA fraction (A, lower panel) and normalized to the ovary.

To address whether the abundance of specific tRNAs matches the actual frequency of the corresponding codons in the transcriptome, we performed RNA-seq of the same zebrafish samples analysed by tRAM-seq, and calculated the frequency of each codon across the mRNA transcriptome (Supplementary Figure S4). Comparing the codon frequency and the tRNA cluster abundance (Figure 3) we found no obvious correlation. Only in the case of the tRNA cluster Lys-CTT, which is highly enriched toward 24 hpf (Figure 3A), we found a consistent increase in the frequency of AAG codons at later time points (Supplementary Figure S4). In contrast, for nearly all tRNAs their different abundance during embryo development appeared to do not correlate with the frequency of the corresponding codons.

### tRNA modification landscape of the developing zebrafish embryo

RNA modifications at the base moiety of nucleoside residues can interfere with RT, causing characteristic “errors” in NGS data (27,35). To detect tRNA modifications that induce RT-errors, we analysed the NGS results of the “mock” samples, i.e. libraries prepared from tRNA of zebrafish samples subject to neither enzymatic, nor chemical treatment to alter modifications before library preparation (Figure 1). In Figure 5, we show the complete landscape of tRNA modifications that we detected in the early developing zebrafish embryo (4-cell stage, 1 hpf) as inferred from the rate of mismatch in the reads spanning all 68 tRNA clusters (see also Supplementary Table 2). Based on available tRNA modification databases (8,36,37), we assigned the identity of the modified sites. We detected high error rates, and thus modification signature, of the following known methylation sites: m^1^A9 and m^1^G9 in many nucleo-cytoplasmic tRNA clusters and all mitochondrial tRNA clusters encoding a purine at position 9, m^1^A14 in Phe-GAA, m^3^C20 in elongator Met-CAT, m^2,2^G26 in many nucleo-cytoplasmic and mitochondrial tRNA clusters as well as m^2,2^G27 in Tyr-GTA, m^3^C32 in several nucleo-cytoplasmic clusters and 2 mitochondrial tRNAs, m^1^G37 in many tRNA clusters, m^3^Ce2 (also known as m^3^C47d) in the variable loop of Leu-CAG and all nucleo-cytoplasmic Ser clusters, and m^1^A58 in nearly all tRNA clusters (Figure 5). We conducted the same analysis on the sequencing data derived from the DM libraries, for which the tRNA fraction was pre-treated by demethylation: as anticipated, all sites of modifications interpreted as m^1^A, m^1^G, m^3^C and m^2,2^G lost the modification signature after demethylation treatment (Supplementary Figure S6), confirming the inferred identity.

**Figure 5.**
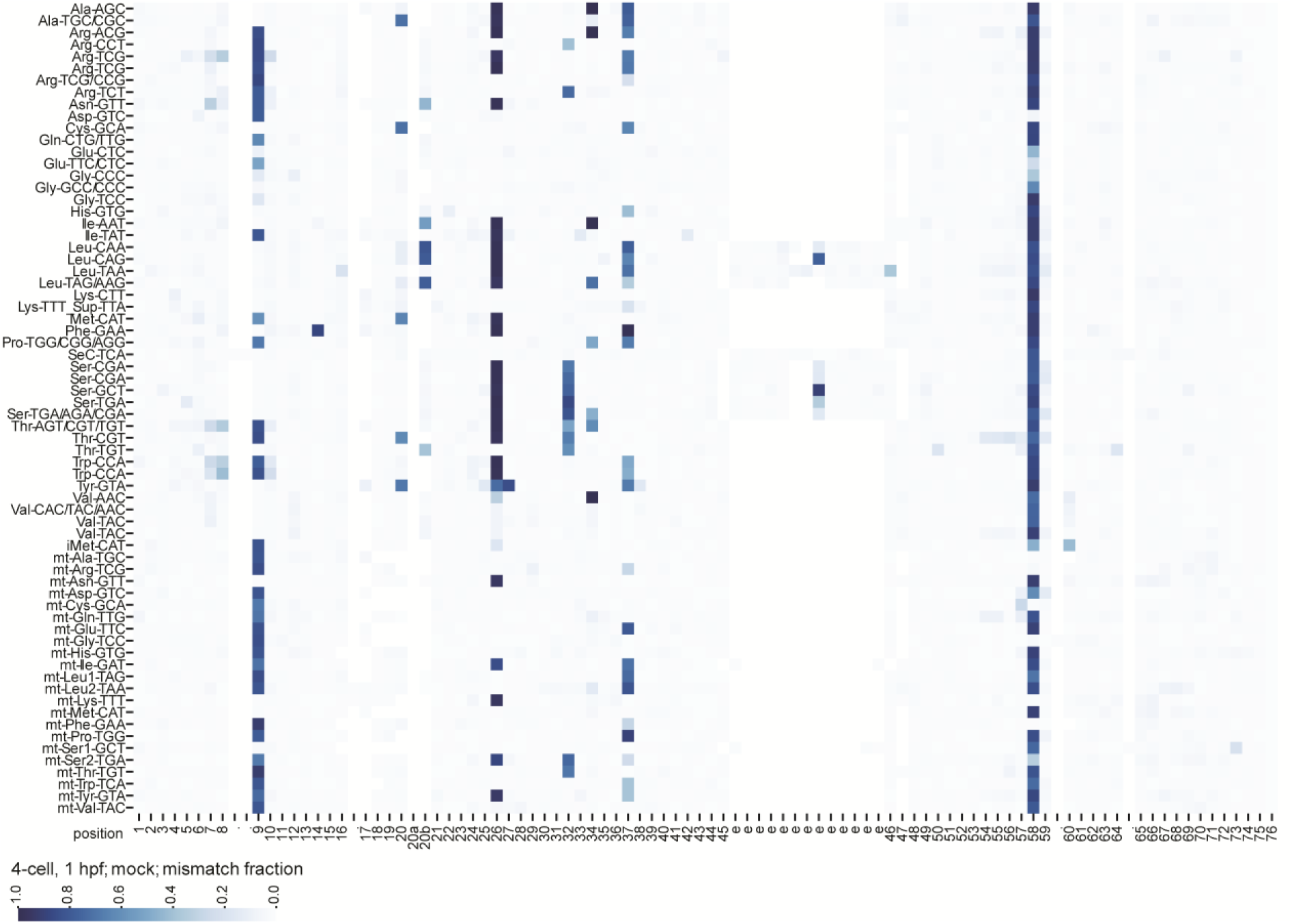
tRNA modification landscape of 4-cell stage zebrafish embryos. Heat-map of mean misincorporation fraction of all nucleo-cytoplasmic tRNA clusters and mt-tRNAs (y-axis). Canonical nucleotide positions are annotated on the x-axis. Nucleotide positions that are rarely present in the tRNA clusters are annotated with a dot. The blue colour scale indicates the mean mismatch fraction across three biological replicates.

We also detected a signature of A-to-I editing. Inosine is produced by adenosine deamination, and it base-pairs with cytosine during RT, causing incorporation of G in reads obtained by Sanger sequencing or NGS. At canonical position 34 of several tRNA clusters, we found prevalent guanosine incorporation in the reads in correspondence of an encoded A34 (Figure 5 and Supplementary File 2). As expected, the mismatch at A34 was not altered by demethylation in the DM samples (Supplementary Figure S6), supporting the interpretation of the site as a case of editing. Another site of A-to-I editing is A37 in tRNA-Ala. In the mock samples, we detected a modification signature in Ala-AGC and Ala-TGC/CGC reminiscent of purine methylation, consisting of a combination of different mismatches (Supplementary File 2). In the DM libraries, the high error rate at position A37 of both tRNA-Ala clusters was still present (Supplementary Figure S6), but converted to nearly exclusively guanosine incorporation (Supplementary File 3), confirming that the original modification found at position 37 of both tRNA-Ala clusters is 1-methylinosine (m^1^I).

In addition to simple base methylations and A-to-I editing, we detected additional modifications interfering with RT. One such RT-interfering modification is the bulky guanosine modification peroxywybutosine (o2yW), which is exclusively found at position 37 of tRNA-Phe in eukaryotes (8) and is expected to cause a major roadblock for RT. Indeed, at position 37 of cluster Phe-GAA we found a high error rate as well as a high fraction of read stops, indicative of prematurely terminated RT at the site (Figure 5 and Supplementary Figure S7). Furthermore, we detected both mismatches and stops at position A37 of mt-Ser2-TGA and mt-Trp-TCA, where the modification 2-methylthio-*N*^6^-isopentenyladenosine (ms^2^i^6^A) is expected.

The region spanning canonical positions 20, 20a, and 20b in the D-loop showed modification signatures seemingly resistant to demethylation in several nucleo-cytoplasmic tRNA clusters (Figure 5 and Supplementary Figure S6), mainly consisting of misincorporations and to a lesser extent RT stops (Supplementary Figure S7). Modifications known to be present in eukaryotes at these sites are 3-(3-amino-3-carboxypropyl)uridine (acp^3^U), pseudouridine (ψ), dihydrouridine (D), and combinations thereof (8,36,37). Besides in the D-loop, ψ is frequently found in the aminoacyl-acceptor stem, D-stem, anticodon stem-loop, variable loop, T-stem, and nearly in all tRNAs at canonical position 55; however, in none of those sites we observed modification signature. Similarly, D is often found at canonical positions 16, 17, and 47, but none of those positions showed modification signature in our tRAM-seq data either. Thus, we suggest that the modifications detected at positions 20, 20a and 20b correspond to acp^3^ sites, being acp^3^U, acp^3^D, or acp^3^ψ.

Another tRNA modification expected to interfere with RT is 2-methylthio-*N*^6^-threonylcarbamoyladenosine (ms^2^t^6^A), to date reported at position 37 of tRNA-Lys-TTT in bacteria and eukaryotes (8). In our clustering pipeline, tRNA-Lys-TTT clustered together with the suppressor tRNA-Sup-TTA. In the cluster Lys-TTT_Sup-TTA, we detected mainly RT stops (Supplementary Figure S6) unaffected by demethylation (Supplementary File 3), with reads terminating at position 38 and thus consistent with the presence of ms^2^t^6^A at position 37.

No modification signature (in the form of mismatches or RT stops) was detected at sites corresponding to known m^2^G, or consecutive dihydrouridine (DD) sites, which have been previously reported to mildly interfere with RT (27,38); this observation suggests that the highly processive reverse-transcriptase that we used, TGIRT, is not affected by these modifications, at least under the experimental conditions that we used.

### tRNA m^5^C methylome of the developing zebrafish embryo

The *C*^5^-methylation of cytosine (m^5^C) is a widely conserved modification found in virtually all RNA classes in the three domains of life, and in higher eukaryotes is introduced by the methyltransferases of the NSUN family, and DNMT2 (also known as TRDMT1) (8). The methylation m^5^C was shown to be important for RNA folding and metabolism, and alterations in the expression/function of m^5^C-installing methyltransferases was linked to developmental defects and cancer (reviewed in (39) and (10)), attracting broad interest. However, previous studies on m^5^C methylation in embryo development have focused on mRNAs (40,41), and the elucidation of the tRNA m^5^C methylome, in particular in vertebrate embryo development, remained unexplored.

Using the BS libraries, we compiled the complete m^5^C methylome of nucleo-cytoplasmic and mitochondrial tRNAs in zebrafish (Figure 6 and Supplementary Table 2). We clearly detected C retention (indicative of m^5^C methylation) in the region spanning canonical positions 48, 49, and 50 of many nucleo-cytoplasmic tRNA clusters, as well as in mt-Leu1-TAG, mt-Leu2-TAA, mt-Thr-TGT, and mt-Tyr-GTA. These positions, located at the junction between the variable loop and the T-stem, are known to be targets of the methyltransferase NSUN2, which has dual localization (nuclear and mitochondrial) (42,43). NSUN2 also methylates C34 of nucleo-cytoplasmic tRNA-Leu-CAA, which is then further modified to 2′-*0*-methyl-5-formylcytidine (f^5^Cm) (44). After bisulfite treatment, f^5^Cm is not expected to be detected as C retention in NGS data like m^5^C, unless an additional protection or reduction step is included in the protocol (45). In the BS libraries, we detected modification signature in the form of C retention at C34 of the cluster Leu-CAA (Figure 6, Supplementary File 4), suggesting that in zebrafish this site is either only partially formylated, or is further reduced to 5-hydroxymethylcytosine (hm^5^C), which behaves like m^5^C in bisulfite treatment and sequencing (46). Similarly, C34 in mt-Met-CAT is expected to be methylated to m^5^C by NSUN3 and then further modified to f^5^C, although the extent of formylation in human cells was subject to dispute (reviewed in (47)). We clearly detected C retention at C34 of mt-Met-CAT (Figure 6), again suggesting that either the site is only partially formylated, or it is further reduced to hm^5^C.

**Figure 6.**
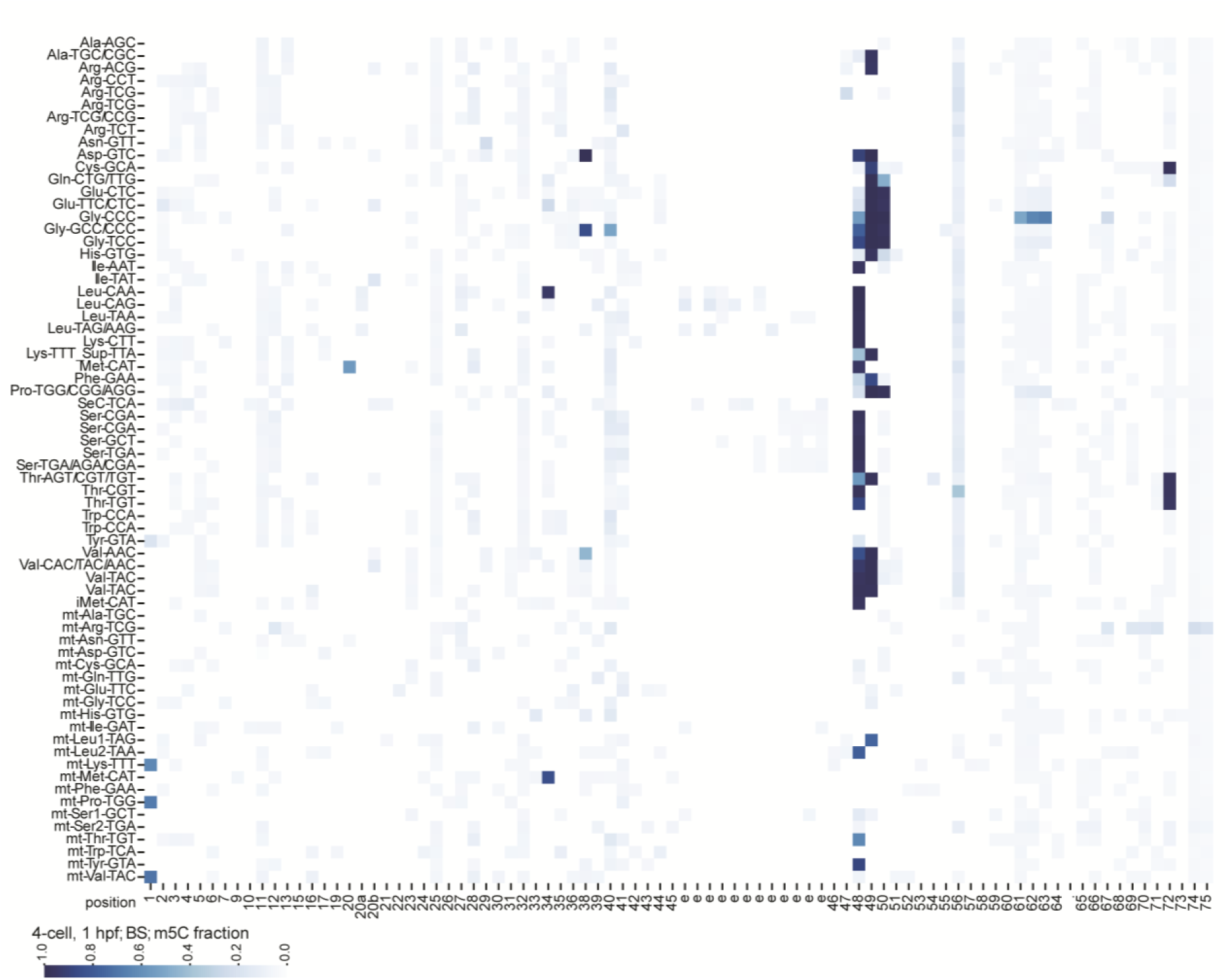
tRNA m^5^C methylation landscape of 4-cell stage zebrafish embryos. Heat-map of C retention, indicative for m^5^C methylation, for all nucleo-cytoplasmic tRNA clusters and mt-tRNAs (y-axis). Nucleotide positions that are rarely present in the tRNA clusters are annotated with a dot. The blue colour scale indicates the mean C retention fraction across two biological replicates. Some tRNA positions show C retention while most likely are not m^5^C methylated: the cluster MetCAT includes tRNAs that either have a C or a U at position 20. The tRNAs possessing a U20 are most likely modified to acp^3^U, which in our RT frequently leads to misincorporation of a G. This results in a C being mapped to a position that has a C as alternative reference, hence the C retention signal. Other likely artefacts are position 61, 62, 63 of the cluster GlyCCC that most likely are the result of incomplete bisulfite conversion due to structure. Lastly, the C retention signal at the 5’ end of the clusters mt-LysTTT, mt-ProTGG and mt-ValTAC are most likely caused by over-trimming of leading Ts (see experimental details in Supplementary File 1).

Finally, we detected signature of m^5^C methylation at C72 of Cys-GCA and all Thr clusters (Figure 6), which are known targets of NSUN6 (48), and at C38 of Asp-GTC and Gly-GCC/CCC, and to a lower extent of Val-AAC/TAC, which are expected to be methylated by DNMT2 (49–51).

### tRAM-seq identifies unexpected tRNA modifications in zebrafish tRNAs

Analysing the tRNA modification landscape in zebrafish, we found unanticipated differences compared to previously described modifications in eukaryotes. In particular, in humans ms^2^i^6^A was reported at position 37 of mt-Phe-GAA and mt-Tyr-GTA; however, in *Danio rerio* both tRNAs encode a guanosine at position 37 (2). Using tRAM-seq, we detected misincorporation at both sites, which was abolished by demethylation (Figure 5 and Supplementary Figure S6), suggesting that in zebrafish G37 of mt-Phe-GAA and mt-Tyr-GTA is rather modified to m^1^G.

In another instance, we detected signature of m^5^C at position 40 of GlyGCC/CCC, although only to an extent of about 50% (Figure 6). To the best of our knowledge, m^5^C40 was previously reported exclusively in archaea and yeast (10), thus we present first evidence of the existence of this modification in a higher eukaryote.

Finally, we observed a misincorporation signature at a U in the variable loop (position e2/47) in the two Ser-CGA clusters (Figure 5 and Supplementary File 2), which disappeared after demethylation (Supplementary Figure S6 and Supplementary File 3). The only known base-methylation of the Watson-Crick face of U is 3-methyluracil (m^3^U), exclusively reported in rRNA (10). Bacterial AlkB can demethylate 3-methylthymine in DNA (52), and the human homolog of AlkB, FTO, was shown to demethylate m^3^U in ssRNA (53). Thus, we can speculate that bacterial AlkB, or one of the mutant variants we used, may be able to demethylate an m^3^U in tRNAs. However, whether tRNA-Ser-CGA in zebrafish really contains a m^3^U in the variable loop will require further validation.

Overall, with our tRAM-seq analysis we elucidated the comprehensive modification landscape of a large array of tRNA modifications in the developing zebrafish embryo, highlighting similarities and differences with tRNA modifications described to date in subsets of tRNAs derived from various organisms.

### tRNA modification is dynamic during embryo development

Comparing the modification signature in tRNAs in the different samples collected (ovary, activated eggs, embryos 1 hpf, 3 hpf, 5 hpf, 10 hpf, and 24 hpf) and processed as “mock”, we observed diverse modification profiles, suggesting dynamic changes during development. In Figure 7 we show selected, representative tRNA clusters and the extent of modification signature at different sites. Most modification sites in tRNA clusters appeared to be constitutively modified, as shown by a stable rate of misincorporation signature over the samples/time-course analysed, as exemplified by m^2,2^G26 in Ala-TGC/CGC, Leu-CAG, Phe-GAA, m^1^A58 in Leu-CAG and Phe-GAA, m^1^I37 in Ala-TGC/CGC, but also evident for many other modifications in different clusters (Figure 7 and Supplementary File 6). It is worth noting that all modification sites in mitochondrial tRNAs appeared to be stable, as inferred from the rate of RT error (Supplementary File 6). However, a subset of modifications in nucleo-cytoplasmic tRNAs appeared to change during embryo development. For instance, misincorporation at position 20 in Ala-TGC/CGC, inferred as a site of acp^3^ modification (see previous section), appeared to be lowest in the ovary samples, nearly double in activated eggs, stable at 1 hpf and 3 hpf, and again decreasing from 5 hpf onward (Figure 7A). We recurrently observed a profile consisting of highest modification signature in activated eggs and the earliest embryonal stages analysed, namely for m^1^A58 in Ala-TGC/CGC and Glu-CTC, m^1^A9 in Asp-GTC, and m^1^A14 in Phe-GAA (Figure 7), but also for m^2,2^G26 in Arg-TCG, m^1^G9 in Glu-TTC/CTC, m^1^A58 in Gly-CCC, acp^3^U20 in Leu-TAG/AAG, m^1^G9 in Met-CAT, acp^3^U20 in Thr-CGT and Thr-TGT, m^2,2^G27 in Tyr-GTA, and to a lesser extent m^3^Ce2 in Ser-CGA (Supplementary File 6). Overall, the ovary samples showed a distinct modification profile, not consistently aligning with either eggs and early embryo, nor with the later stages analysed (Figure 7 and Supplementary File 6). This might reflect the content of oocyte precursors at different stages of maturation that may differ substantially from mature eggs, and/or their combination with the proportion of maternal (mature) tissue content.

**Figure 7.**
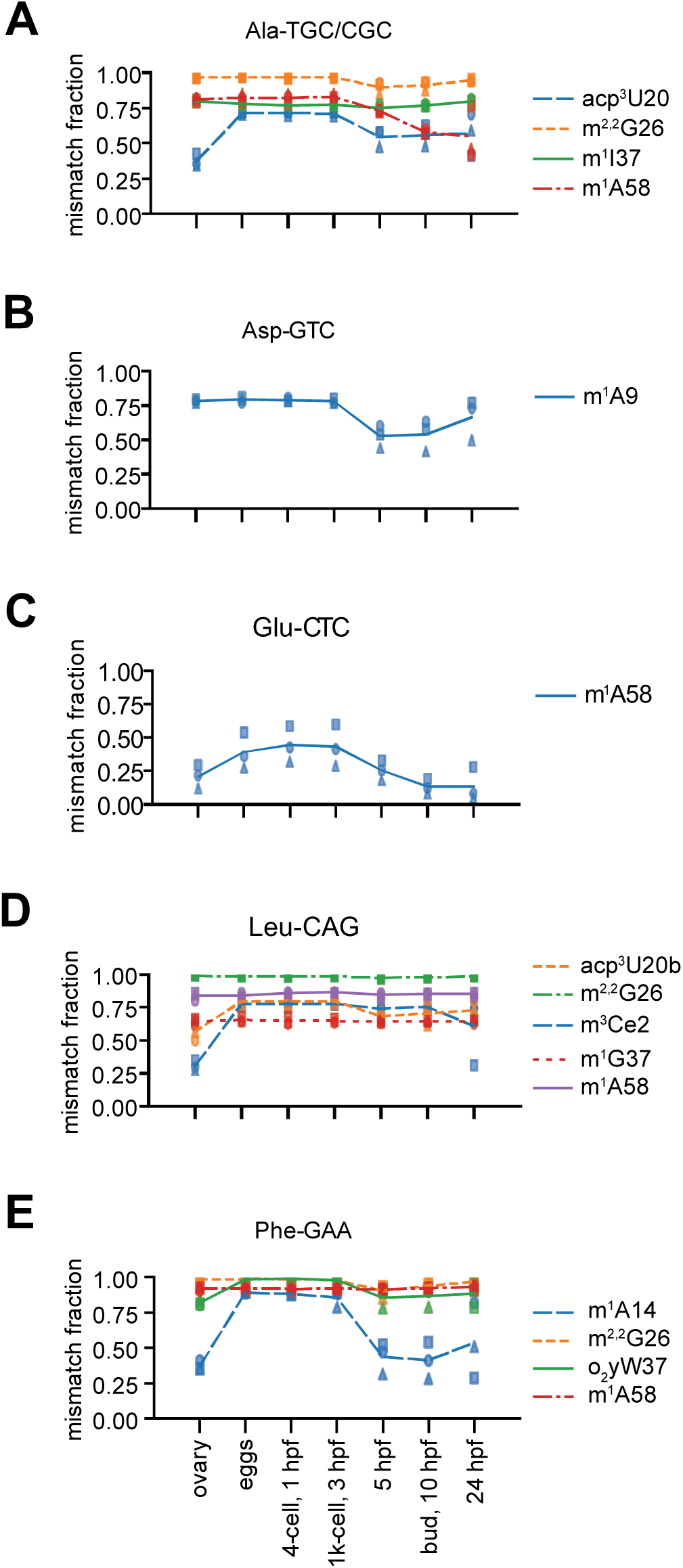
tRNA modification dynamics during zebrafish embryo development. (**A-E**) Misincorporation fraction in five selected tRNA clusters throughout developmental stages of zebrafish embryos, plus ovary (x-axis). Graphs are shown for tRNA nucleotide positions that have a misincorporation fraction equal or greater than 0.15 in at least one time point. The tRNA clusters are indicated on top of the plots; the predicted modifications causing the misincorporation are indicated in the legend on the right. Lines represent the mean of the three biological replicates, which are indicated by square, circle and triangle.

We validated the observed tRNA modification dynamics for two representative modification sites by primer extension (27). For both cases, m^1^A14 in Phe-GAA and m^1^A58 in Glu-CTC, the modification profile obtained by primer extension closely resembled the one observed by tRAM-seq, confirming the reproducibility of our results and the observed dynamics of tRNA methylation during embryo development (Figure 7 and 8).

**Figure 8.**
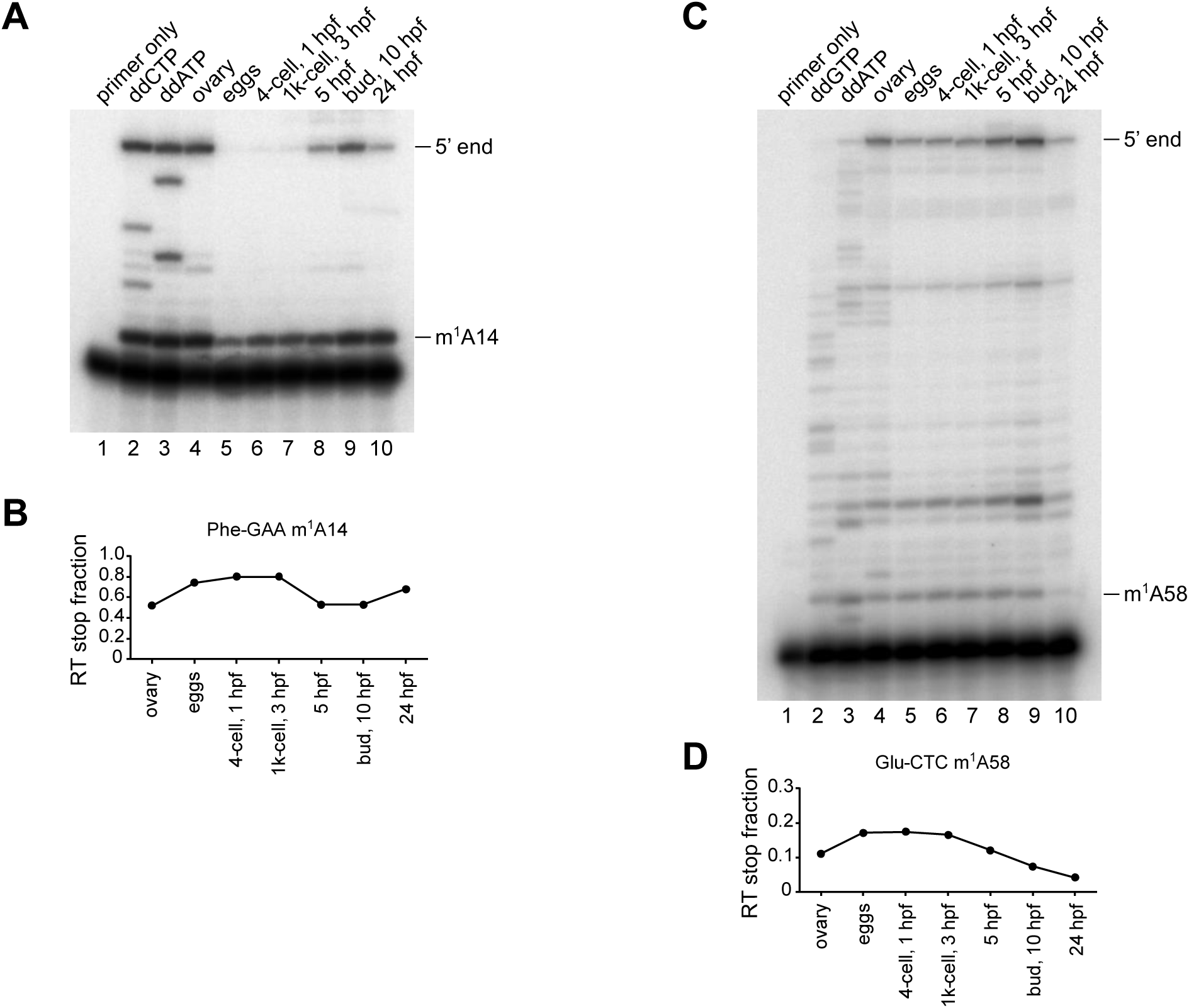
tRNA modification detection by primer extension. Primer extension analysis of (**A**) m^1^A14 in PheGAA and (**C**) m^1^A58 in GluCTC performed using AMV RT on total RNA extracted from zebrafish samples and a corresponding radiolabelled primer. In lane 1 only the primer was loaded; in lanes 2 and 3 RT products that were reverse transcribed from zebrafish ovary total RNA in presence of the indicated ddNTPs were loaded. Lanes 4-10 contain the RT products reverse transcribed from each sample. RT stops caused by modification and the full-length cDNA are annotated on the right side of the image. (**B**) and (**D)**: quantification of m^1^A14 in PheGAA and m^1^A58 in GluCTC detected as RT stops in (A) and (C), respectively. The RT stop fraction is quantified by dividing the modification-induced RT stop by all RT stops in the lane (excluding the unextended primer).

Also in the case of m^5^C modifications detected by bisulfite treatment, we observed dynamics across the analysed samples. While most sites displayed stable C retention, thus m^5^C modification extent, a subset of sites showed a modulated profile. Figure 9 shows the m^5^C profile of selected, representative tRNA clusters. Remarkably, all instances of m^5^C sites apparently modulated, i.e. whose extent of C retention changed during zebrafish embryo development, were at positions 40, 48, 49, and 50, which are all targets of the methyltransferase NSUN2 (Figure 9 and Supplementary File 7). However, not all NSUN2 target sites were modulated, and even adjacent target sites of NSUN2 within the same cluster were differentially modified, for instance in the case of positions 49 and 50 of Gln-CTG/TTG, and 48 and 49 of Phe-GAA (Figure 9B and E).

**Figure 9.**
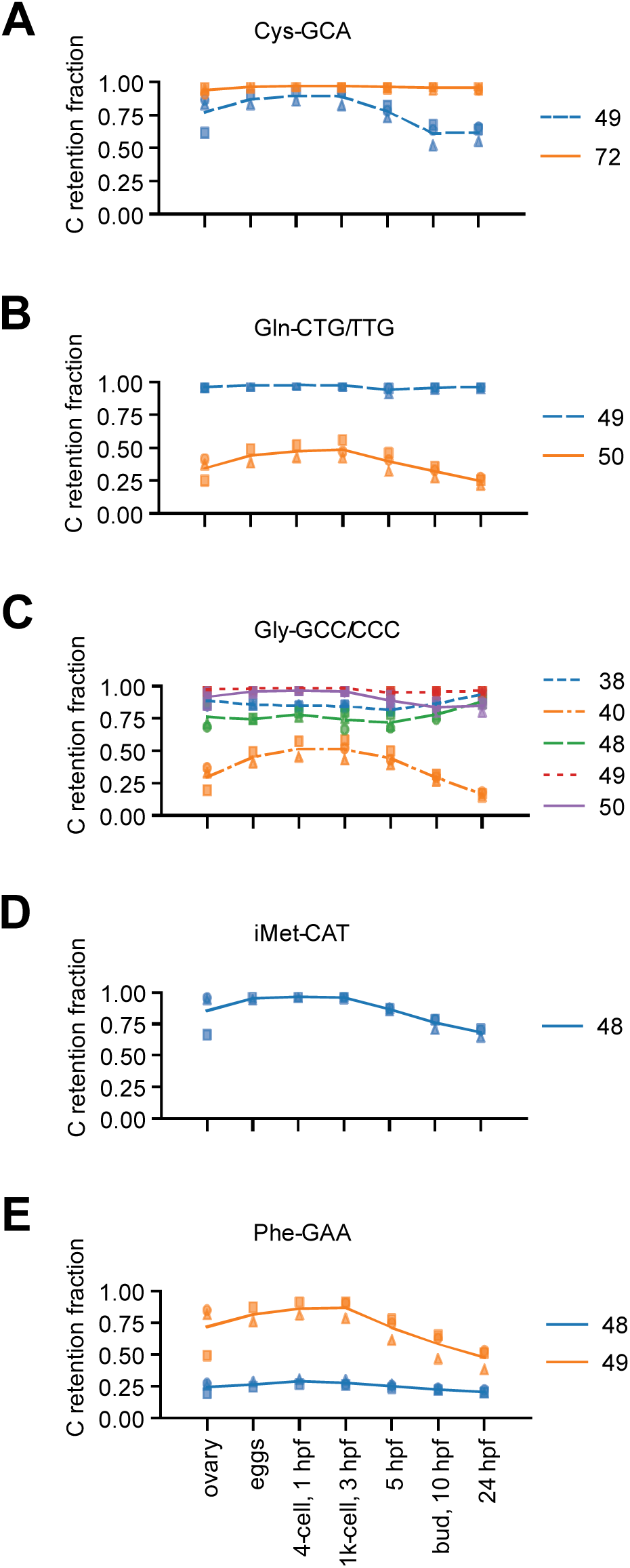
tRNA m^5^C methylation dynamics during zebrafish embryo development. (**A-E**) C-retention fraction, indicative of m^5^C modification, of five selected tRNA clusters throughout developmental stages of zebrafish embryos, plus ovary (x-axis). Lines show the mean C-retention fraction of positions that have a C-retention fraction equal or greater than 0.15 in at least one time point (n=3, except for activated eggs and 4-cell stage where n=2).

To determine whether the expression of tRNA modification enzymes also changes during zebrafish embryo development, we referred to the list of all tRNA modification enzymes identified to date (54), and analysed their expression profile in our gene expression data sets (see first results section and Supplementary Figure S8). The steady-state levels of the mRNAs of many tRNA modification enzymes appeared to be changing over the time course analysed, with some highly expressed in eggs and early zebrafish embryo, and others showing higher expression levels at later time points (Supplementary Figure S8). However, the mRNA levels of modification enzymes did not appear to generally correlate with the abundance of the corresponding modification. For instance, the mRNA of the zebrafish homologue of TRMT5 appears to be more abundant in eggs, 1 hpf and 3 hpf, with levels progressively declining at 5 hpf, 10 hpf and 24 hpf (Supplementary Figure S8). TRMT5 is responsible for m^1^G37 and m^1^I37 methylation in many nucleo-cytoplasmic and mitochondrial tRNAs (8,55). According to our tRAM-seq analysis, m^1^G37 and m^1^I37 appeared to be substantially stable in all tRNA clusters where they were detected. In the case of TRMT10B, responsible for m^1^A9 in Asp-GTC (34), we observed a slight, transient decrease of mRNA levels from 1 hpf to 10 hpf, and subsequent increase at 24 hpf (Supplementary Figure 8). This profile is tentatively compatible with the decrease in m^1^A9 in Asp-GTC around 5-10 hpf and subsequent new increase at 24 hpf (Figure 7B). The mRNA steady state levels of TRMT10A, responsible for m^1^G in many nucleo-cytoplasmic tRNAs (34), progressively declined from activated eggs until 5-10 hpf (Supplementary Figure S8); however, the modification signature of m^1^G9 was substantially stable for all tRNAs target of TRMT10A, with the exception of Glu-TTC/CTC and Met-CAT, where it apparently decreased from 5 hpf onward (Supplementary File 6). Also, the mRNA of CDKAL1, responsible for the modification of t^6^A (silent in our tRAM-seq modification detection) to ms^2^t^6^A37 in tRNA Lys-TTT (56) appeared to increase over time from activated eggs to 24 hpf; however, the modification signature for ms^2^t^6^A37 that we observed in the cluster Lys-TTT_Sup-TTA, consisting mainly of RT stops at position 37, was substantially unchanged (Supplementary Figure S9). Similarly, the modification profile of further tRNA modifications did not match the apparent expression levels of the corresponding tRNA modification enzymes, as inferred from their mRNA levels.

In conclusion, here we show for the first time that tRNA modifications are dynamic during vertebrate embryo development. The molecular mechanism and determinants of the specific modification patterns cannot be attributed to the apparent expression levels of modification enzymes, and can entail more complex mechanisms affecting tRNA substrate-selection and base-selection specificity.

## DISCUSSION

The apparent redundancy of tRNA genes is known since decades, and the expansion of the tRNA genes’ copy number was suggested to be linked to the frequency of codons in the mRNAs of abundant proteins, at least in bacteria and yeast (57,58). The investigation of the actual (eventually differential) expression levels of tRNAs used to be limited to tedious hybridization-based methods like microarray and northern blotting. Nevertheless, early work showed that in higher eukaryotes the pool of expressed tRNAs can vary between tissues, proliferative states, and between physiological and pathological conditions like cancer (59,60).

Here we built upon recent developments in NGS-based methods for the study of tRNAs, and devised our wet lab and computational analysis pipeline, tRAM-seq, to investigate the dynamics of tRNA expression and modification in zebrafish embryo development. We combined a previously published protocol for tRNA sequencing using the highly processive RT TGIRT with improved reaction conditions (11) with the incorporation of randomized ligation termini aimed to minimize ligation bias (23), which we also used for PCR product deduplication. Also, similar to others (12,13,61), we used the demetylase AlkB and its mutant variants to demethylate our DM samples specifically for tRNA abundance analysis, aiming to minimize any bias in representation of heavily-versus lightly-modified tRNAs. Noteworthy, we obtained sufficient coverage also for tRNAs carrying bulky modifications that are known to pose a hard stop to RT and that are not removed by AlkB, namely o^2^yW37 in Phe-GAA and ms^2^t^6^A37 in Lys-TTT (8). Furthermore, the comparison between the mock and DM samples allowed us to substantiate the inferred type of modification detected at specific positions.

Unique to our analysis pipeline was the use of the actual sequencing data to select and construct the best-matched reference to perform the final mapping of the tRNA reads. This approach proved effective for the analysis of *D. rerio* NGS data, which was not addressed by previous work probably due to the extreme expansion of tRNA genes’ copy number in zebrafish (over 20000 including low-scoring predictions) (2). We further collapsed the still large number of tRNA genes found expressed in the DM samples into representative clusters, based on both sequence similarity and actual fraction of multimapping reads. The sequence differences in the clustered references are visualized in our coverage plots, adding a layer of information available in the analysis. Moreover, our mapping strategy did not make use of “masking” known modified positions to accommodate mismatches at those sites (11), rendering our analysis independent of previous knowledge of modifications (which may be scarce for organisms other than the most investigated models).

Using tRAM-seq we elucidated for the first time the dynamics of tRNA expression and modification during the embryo development in the vertebrate model zebrafish. Interestingly, Asp-GTC appeared to be the most abundant tRNA, in particular at 10 hpf and 24 hpf, with up to 10% of the total tRNA reads mapping to the cluster Asp-GTC (Figure 3A and B). Asp-GTC also displayed a strong modulation across the samples analysed, increasing from about 5% up to 10% of the total tRNA reads. The only previous study that quantified tRNA expression in zebrafish embryo, at the single time point of 6 hpf, similarly found Asp-GTC being the most abundant tRNA (62); however, the authors did not observe a high frequency of aspartate-encoding codons in the corresponding transcriptome. Likewise, we did not observe a correspondence between the high abundance of tRNA-Asp and the frequency of aspartate-encoding codons in our transcriptomic data (Figure 3 and Supplementary Figure S5). Another abundant tRNA in our datasets was Lys-CTT, which also showed modulation during zebrafish embryo development: its abundance levels, in terms of fraction of total tRNA, nearly doubled from about 3% in activated eggs and 4-cell stage, to about 6% at 24 hpf (Figure 3). Lys-CTT was the only case where we observed a correspondence between tRNA abundance and frequency of codons, as the frequency of the codon Lys-AAG similarly increased over the time course analysed (Supplementary Figure S5), possibly suggesting that the increase in tRNA-Lys-CTT is needed to match the translational requirements of Lys-containing proteins. Still, for the large majority of the tRNA clusters we did not observe a consistent correspondence with the matched codon frequency, suggesting that the levels of the different tRNAs may not be directly linked to the overall coding exome. Interestingly, it was previously suggested that the selection of non-optimal (rare) codons may play a role in coordinating the expression of specific subsets of genes (63,64). Thus, the actual levels of tRNAs may have a more complex impact on the rate of translation of different genes, and the observed changes may have evolved to support the very specific, rapidly changing expression program of the developing embryo. Alternatively, the relative abundance of different tRNAs may be linked to alternative tRNA functions, for instance in interaction with proteins, RNAs, or metabolic pathways (1). In the case of mitochondrial tRNAs, the apparent higher levels of mt-Val-TAC may be due its incorporation as a subunit of the mitochondrial ribosome (65), and the observed increase from 5 hpf may be due to the new synthesis of organellar ribosomes.

When we dissected the tRNA clusters into the individual isodecoders, we observed a major switch in prevalent tRNAs present in activated eggs and early embryos versus embryos at later stages. The switch in tRNA expression appears to happen from 5 hpf onward, which corresponds to the start of gastrulation, i.e. the massive migration of embryonal cells and establishing of distinct embryonal layers (18). The observed change in tRNA ensemble can be reasonably attributed to the replacement of the original, maternally deposited tRNAs present in the egg by zygote-expressed tRNAs. Whether the maternal tRNAs decline is due to dilution by newly transcribed tRNAs, regular turnover, or active/selective degradation remains to be clarified. In any case, how the different tRNA genes scattered across the zebrafish genome are selectively activated for transcription is not known. The eukaryotic RNA polymerase responsible for tRNA transcription is Pol III, and it was recently shown that Pol III transcribes different tRNA loci in human induced pluripotent stem cells versus cardiomyocytes and neurons (66). Still, how different tRNAs can be expressed at different stages of embryo development remains to be clarified.

Similar to the analysis of tRNA expression, the study of (t)RNA modifications has lagged behind due to technical challenges. Early work relied on the use of radioactive labelling and separation by TLC (67), or the use of high-performance liquid chromatography and mass spectrometry (68). These methods require the isolation of large amounts of the specific RNA of interest to be analysed, and, as a consequence, are severely limited in applicability to endogenous RNAs. The recently available NGS-based methods overcome this limitation allowing to investigate a broad range of modifications at transcriptomic level with relatively little amount of starting material (16). These new methods have thus opened the avenue for investigating the differential modification of tRNAs across tissues/samples, conditions, developmental stages, etc. It is worth noting that whilst A-to-I editing is consistently read as G in sequencing experiments, other modifications are detected as a combination of different misincorporations/stops, where also the correct base is incorporated to a certain extent during RT. The modification signatures observed, consisting of misincorporations and RT stops, cannot be directly translated into absolute values since the type and extent of RT errors depends on multiple factors, including the type of modification itself, the adjacent sequence, and the RT used (11,30,69). However, the percentage of error represents the minimum possible extent of modification, and it is reasonable to assume that high misincorporation rates correspond to complete or nearly complete modification. In any case, the relative comparison of individual modification sites between matched samples and/or time points (like in our case) remains valid as the type of modification and sequence context remain unaltered.

To date, the only studies of tRNAs and their modification profile during development were conducted in the amoeba *Dictyostelium discoideum*, which develops from single cell to a multicellular fruiting body in response to starvation (70), and in *Drosophila* (71), whilst no study addressed so far tRNA expression and modification during development in vertebrates. Here we profiled eleven different tRNA modifications, namely: I, m^1^I, m^1^G, m^1^A, m^2^_2_G, m^3^C, m^5^C, acp^3^U, ms^2^i^6^A, ms^2^t^6^A, and o^2^yW. We showed that the extent of modification at a subset of tRNA positions changes during zebrafish embryo development, prevalently displaying higher modification signature in activated eggs and early developmental stages, and a decrease in modification after 5 hpf. These dynamics are unlikely to be generally due to changes in the abundance of the responsible modification enzymes, as we did not observe a correlation with their mRNA expression levels (Supplementary Figure S8) or with their protein levels in similar samples that we recently reported (72). Furthermore, different sites modified by the same enzyme showed different modification profiles. In a few instances, changes of modification signature could be attributed to an actual change in abundance of different tRNAs within a cluster, of which only some can be modified. This is reasonably the case for I34 in Thr-AGT/CGT/TGT and Val-CAC/TAC/AAC, which are the only sites where we rather observed an increase in modification signature at 10 hpf and 24 hpf as compared to earlier stages (Supplementary File 6). Here the increase in modification signature is due to the relative increase of abundance of the Thr-AGT and Val-AAC isodecoders within the respective cluster (Figure 3C and Supplementary File 5). In other instances, the different modification extent may be the consequence of finer differences in the activity and/or affinity of the modification enzyme for specific tRNA positions. For example, in the case of the cluster Ala-TGC/CGC, the isodecoder Ala-TGC-1 is prevalently expressed in early time points, and after 5 hpf appears to be replaced by Ala-TGC-4; in parallel, the overall extent of m^1^A58 methylation in the cluster appears to decline. Actually, the extent of m^1^A58 modification remains consistently high for Ala-TGC-1 and low for Ala-TGC-4, thus the apparent lower modification of the cluster at later time points is due to the change in relative abundance of the different isodecoders. Ala-TGC-1 and Ala-TGC-4 differ in the variable loop by having U47 or C47, and in the T-stem where the base-pair 50-64 is U:A or C:G, respectively. Interestingly, the methylation of A58 by the TRMT61A-TRIMT6 complex requires conformational rearrangements of the T- and D-stem loops (73); thus, the identity of the bases and/or base-pairs present may affect the flexibility of the regions involved, and ultimately the efficiency of methylation. Given our observation of a change in isodecoder expression during embryo development, it is intriguing to hypothesize that differences in sequence (and possibly structure as consequence) between different isodecoders may act as determinants for -or affect recognition by-modification enzymes, entailing a coordinated modulation of isodecoder expression and modification. The remodelling of the tRNA transcriptome in terms of isodecoder expression and modification during embryo development holds the potential of fine tuning the function of tRNAs in protein translation and beyond. The full range of functional consequences of this phenomenon remains an open question.

## DATA AVAILABILITY

All sequencing data were deposited in the Sequence Read Archive (SRA) and are available under the accession number PRJNA1061456.

The bioinformatics analysis results are available at FigShare 10.6084/m9.figshare.25035872. An implementation of the tRAM-seq bioinformatic workflow within snakemake is available at FigShare.

## SUPPLEMENTARY DATA

Supplementary Data are available on line.

## FUNDING

This work was supported by the Austrian Science Fund (FWF) [F80 to EV, AP, and ILH][Y 1031-B28 to AP]; and the Novo Nordisk Foundation [NNF21OC0066551]. Research in the lab of AP was funded by the Research Institute of Molecular Pathology (IMP); Boehringer Ingelheim; the European Research Council (ERC) [Consolidator grant 101044495/GaMe]; and the Human Frontier Science Program (HFSP) [Young Investigator Grant RGY0079/2020]. A. Chugunova was supported by the European Commission [MSCA-IF-EF-SE 895790].

Funding for open access charge: Austrian Science Fund [F80].

## ACKNOWLEDGEMENTS

The authors thank Ronald Micura (University of Innsbruck) for providing the m^1^G-modified RNA oligonucleotide, Alan Lambowitz (University of Texas) for providing TGIRT, Fatinah El-Isa for assistance with the preparation of NGS libraries, and the Core Facilities of the Medical University of Vienna, a member of VLSI, for sequencing and initial data analysis.

## AUTHOR CONTRIBUTION

EV conceived the project. EV and AP devised the experiments. TR and AC performed the experiments. EV and TR analysed the results of wet lab experiments. MW and IH devised and conducted the bioinformatic analysis of the NGS experiments. EV and TR wrote the manuscript. All authors revised and approved the final manuscript.

## CONFLICT OF INTEREST

None declared.

## Notes

### Competing Interest Statement

The authors have declared no competing interest.

