## Supplementary File 1 for "tRNA expression and modification landscapes, and their dynamics during zebrafish embryo development"

### SUPPLEMENTARY TEXT

#### tRAM-seq analysis

**Preprocessing**: For each sample, we removed adapters with cutadapt (1) (-q 10 -e 0.15 --minimum-length=29 --discard-untrimmed -a <adapter sequence>) and extracted UMIs with umi-tools (2), trimmed leading Ts (possibly untemplated nucleotides introduced by TGIRT (3)) with cutadapt (-g 'XTTTTTTTTTTTTTTT' -e 0.0001, --minimum-length=20), filtered low-quality reads as well as short reads (<20nts) with fastp (4) ( --length_required=20 --trim_poly_x poly_x_min_len=7 -p --low_complexity_filter --complexity_threshold=30) and performed standard quality control before and after each step (fastp and custom code). On average in this pre-processing step 29% of raw reads were removed with the majority (~24%) being removed in the first size selection steps. This resulted in 496495 to 11737789 filtered reads per sample, on average 3473257 (Supplementary Table 2).

**Reference genome**: As an initial reference set we collected 22 mitochondrial tRNA sequences from mitotRNAdb (5) (http://mttrna.bioinf.uni-leipzig.de/mtDataOutput/ August 2022) and 20577 genomic (nuclear) sequences (including 8673 high scoring genes) from a tRNAscan-SE (6) run as provided in the GtrnaDB (7,8) (gtrnadb.ucsc.edu/genomes/eukaryota/Dreri11/danRer11-tRNAs.tar.gz August 2022), extended the sequences by 3'-CCA and removed duplicates, resulting in a set of 10738 unique sequences.

In pre-studies we found that our NGS data includes reads from tRNA genes that are not covered in the high scoring GtrnaDB set. Therefore, we opted to include all predicted genes (including e.g. low confidence predictions and pseudo genes) and performed a mapping of all demethylated samples on this initial set of tRNA candidate genes with segemehl (-D 1 -M 200 -E 500 --accuracy 85) (9-11). Any gene that had a coverage of at least 500 RPM was included in the reference genome for further analysis. As opposed to later mapping steps, here multimappers were not assigned to one possible reference at random but were assigned to the reference that had the most reads mapping to it. This ensured that references that had no (or next to no) uniquely mapping reads assigned to them got filtered out.

After a trial run of the whole analysis pipeline we further manually refined the reference set. Since DM, BS, and Mock treated samples were sequenced in the same sequencing run, there was a potential for cross contamination due to index hopping, furthermore misincorporation due to a modification could assign reads to a gene that is not actually expressed. We therefore removed references with low coverage that only differ from a higher coverage reference at positions corresponding to a C to T transition, or that have a high misincorporation rate from known modification (e.g. references with G in known inosine positions). In addition, we re-added references that feature A instead of G nucleotides in known inosine positions.

For the mitochondrial tRNAs, we noticed inconsistencies between the sequence of in the database (5) and our own sequencing reads. We proceeded to isolate DNA from zebrafish samples and amplified the regions of mitochondrial DNA spanning mt-Arg, mt-Asp, mt-Ser(UCN) and mt-Thr. In mt-Arg, mt-Asp, and mt-Ser(UCN), we confirmed the SNPs observed in our tRAM-seq data, whilst for Thr we observed two alternative genotypes present in different samples, encoding either G or A at position 57 (Supplementary Figure S3). This diversity may be due to SNPs present in individual zebrafish within the pool of animals used per time point, or alternatively be due to heteroplasmy within individual animals in the pool. In any case, we updated our references to include the observed mitochondrial sequence variants. In conclusion, our reference included a total of 223 unique tRNA reference sequences.

**Mapping**: The mapping was performed with segemehl using short RNA optimized parameters (-E 500, -M 200, -D 3). The required mapping accuracy was adjusted to reflect the different number of modifications and thus different misincorporation frequencies in mock, demethylated and bisulfite treated samples (accuracy 80, 85 and 90 respectively). The mismatch tolerance also allows for mapping of SNPs and possibly left over non-templated nucleotides. For bisulfite treated samples segemehl was used with the additional option '-F 1' for BS mapping (and RNA specific post-processing). All samples achieve a mapping rate above 80% (DM: 80-97%; MOCK: 82-97%; BS: 81-94%). However, on average around 70% of the mapped reads are multimappers (DM: 55-70%; MOCK: 70-87%; BS: 63-79%). After mapping, any duplicated reads were removed based on their UMI. On average 5% of reads were filtered out in this step, with a range from two 2% to 18%.

**Clustering**: Given the high similarity, we decided to resolve the high number of multimappers by clustering similar tRNA genes. To retain as much resolution as possible, we decided to perform a two-step clustering based on sequence similarity and multimapper information. In the first step we align all reference sequences to the RFAM (12-14) tRNA covariance model (RF0005) with cmalign (15) and compute a pairwise edit distance of the aligned sequences. We recursively merge any references/clusters that have an edit distance below 4. For the second step, we first count the number of multimapping reads between two clusters. If more than 50% of reads that can be mapped to references in one cluster are multimappers with a second cluster, the two clusters are merged. This results in 68 well resolved clusters that retain full isoacceptor resolution except for one cluster containing lysine (anticodon TTT) and a TTA suppressor tRNA. In most cases the resolution is even better than isodecoder resolution (see supplement table ‘cluster-composition.tsv’). Less than 0.1% of reads are multimappers between clusters.

**Alignment**: The canonical positions within each reference sequence were assigned based on the alignment of the reference to the RFAM tRNA covariance model as described in the previous paragraph. In the first step, canonical positions according to (16) were manually assigned to corresponding positions in the RF0005 alignment. In a second step canonical positions of the reference sequences were automatically assigned based on the alignment of the reference to the RF0005 model. The following lines represent the consensus sequence and structure of the RF0005 tRNA alignment and the assigned canonical positions.

```
#=GC RF                    GgagauaU.A.GCucAgU...GGU...AgaGCg.u.cgGaC.UuaaAAuCcg.aag........................g...cgcg.GGU.UCg.Aa..UCCcg.c.uaucucC.a
#=GC SS_cons               (((((((,.,.<<<<___...___..._>>>>,.<.<<<<_.______>>>>.>,,........................,...,<<<.<<_.___.__.._>>>>.>.))))))).:
                           00000000 0 11111111 111222 222222 2 22333 3333333444 444-eee-eeeeeeeeeeeee-eee--44  4455 555 555 55  66666 6 6666777 7
                           12345678 9 01234567 789000 123456 7 89012 3456789012 345                        67  8901 234 567 89  01234 5 6789012 3
                                               A   AB
```

**Abundance**: The abundance of each cluster was computed based on the DM samples by mapping to the manually refined reference set, random assignment of multi mapping reads to one of the best matching references, counting of the mapped reads per reference, summing up the abundance of all references in a given cluster and normalizing the abundance count by the total count of mapped reads.

**Misincorporation rates**: Possible modification sites and changes of modification level are identified by computing position-wise misincorporation rates for all samples and all treatments. For each reference position, the total number of reads that cover the given position are counted. Furthermore, we count how many of those reads mismatch at the given position. The overall misincorporation rate per position in the clusters is computed by first adding up the mismatch counts and total coverage count for equivalent positions in the tRNA references within the given cluster and then dividing the mismatch count by the total count. Equivalent positions between tRNA references within a cluster are defined by the covariance model based alignment described in the previous paragraph on the clustering approach.

**RT stop fraction**: In addition to the misincorporation rate, RT stops can indicate modified nucleotides. We assume that if a modification at position n leads to a RT stop, the last nucleotide in the generated cDNA is one position upstream, corresponding to one position downstream (n+1) towards the 3'-end of the tRNA reference sequence. Therefore, the number of reads that end at n+1 was divided by the total number of reads that map to n+1 in the given reference tRNA. Analogous to the misincorporation rate, the overall RT stop fraction of position n within a cluster was computed by summing all read-end counts at n+1 of equivalent positions and dividing by all reads that map to n+1 in the given cluster.

**m$^5$C fractions**: Based on the BS treated samples, we also detect putative m$^5$C modification sites and m$^5$C modification dynamics. Unmodified Cs are read as Ts after successful bisulfite treatment while m$^5$Cs are retained as Cs. Thus, m$^5$C modification calls can be based on a C-retention rate. For single tRNA references, the C -retention rate is only computed for C positions and the count of reads that contain a C in the mapped position is divided by the count of mapped reads that contain a C or a T in the given position.

Bisulfite treatment leads to an increase of multimappers, as any T in a read could originate either from a native T or a bisulfite converted C and could therefore be mapped to either reference sequence. To account for this, we selected to compute the C-retention rate in clusters not just based on reads that map to reference Cs but also count Ts that map to native T positions (since these could also originate from bisulfite converted Cs). Thus, per cluster and per position, we obtain the count of mapped Cs as the number of reads with a C that is mapped to a reference C; the count of mapped Ts is the number of reads with a T that is mapped to either a reference C or T. The C-retention rate is computed as usual as the count of Cs divided by the sum over the counted Cs and Ts. This C-retention rate may yield values lower than the actual m$^5$C methylation level if some of the references contain a native T at the position of interest. This C-retention rate is only representative for those references in a cluster that have exhibit a C or T in the given position, thus it does not provide any information on how representative these C and T containing references are for the full cluster. Furthermore, apparent changes in the C-retention rate can also originate from changes in the abundance ratio between references with native Cs and native Ts. Such cases can be detected by comparing to the per reference abundance analysis in the DM samples.

**Principal components analysis**

To evaluate the reproducibility between replicates and similarities between time points, a principal components analysis was performed on the abundance data. As input data the normalized abundance (in RPM) of each tRNA cluster was used (Supplementary Figure S10). To give each cluster the same weight, the abundance of each cluster was standardized by removing the mean abundance of the cluster over all samples and scaling to unit variance.

Similarly, a principal components analysis was performed on the vst data of the mRNA-seq data (Supplementary Figure S11).

**Supplementary Table 1 Cluster composition.**

Composition of the tRNA clusters in terms of tRNA gene reference identity and relative contribution per anticodon (based on anticodon coverage). Mitochondrial tRNAs with SNPs identified in this study were added to the reference, and are indicated with suffix “s”.

| cluster name | cluster ID | anticodon ratio | tRNA name(s) |
|---|---|---|---|
| Ala-AGC | 58 | 1 | Ala-AGC-2; Ala-AGC-3; Ala-NNN-10; Ala-NNN-11; Ala-NNN-3 |
| Ala-TGC/CGC | 59 | 0.78_0.22 | Ala-CGC-2; Ala-CGC-3; Ala-TGC-1; Ala-TGC-4 |
| Arg-ACG | 28 | 1 | Arg-ACG-8; Arg-ACG-6; Arg-ACG-5; Arg-ACG-4; Arg-ACG-3; Arg-ACG-1 |
| Arg-CCT | 42 | 1 | Arg-CCT-2; Arg-NNN-27 |
| Arg-TCG | 6 | 1 | Arg-TCG-3 |
| Arg-TCG | 33 | 1 | Arg-TCG-2; Arg-TCG-1 |
| Arg-TCG/CCG | 63 | 0.54_0.46 | Arg-CCG-2; Arg-CCG-3; Arg-CCG-5; Arg-TCG-8; Arg-NNN-115; Arg-NNN-68 |
| Arg-TCT | 54 | 1 | Arg-NNN-13; Arg-TCT-1; Arg-TCT-12; Arg-TCT-2; Arg-TCT-20; Arg-TCT-21; Arg-TCT-5; Arg-TCT-6 |
| Asn-GTT | 47 | 1 | Asn-GTT-65; Asn-GTT-5; Asn-GTT-4; Asn-GTT-3; Asn-GTT-2; Asn-GTT-16; Asn-GTT-15; Asn-GTT-29 |
| Asp-GTC | 67 | 1 | Asp-GTC-1; Asp-GTC-2; Asp-GTC-4; Asp-GTC-6 |
| Cys-GCA | 32 | 1 | Cys-GCA-1; Cys-GCA-2 |
| Gln-CTG/TTG | 60 | 0.69_0.31 | Gln-CTG-1; Gln-CTG-2; Gln-CTG-3; Gln-CTG-4; Gln-TTG-6; Gln-TTG-4; Gln-TTG-3; Gln-TTG-1 |
| Glu-CTC | 49 | 1 | Glu-CTC-1; Glu-CTC-23 |
| Glu-TTC/CTC | 40 | 0.87_0.13 | Glu-CTC-17; Glu-TTC-12; Glu-TTC-2; Glu-TTC-37; Glu-TTC-4; Glu-TTC-49 |
| Gly-CCC | 16 | 1 | Gly-CCC-20 |
| Gly-GCC/CCC | 52 | 0.89_0.11 | Gly-CCC-4; Gly-CCC-1; Gly-CCC-2; Gly-GCC-10; Gly-GCC-2; Gly-GCC-20; Gly-GCC-3; Gly-GCC-6; Gly-GCC-7 |
| Gly-TCC | 44 | 1 | Gly-TCC-2; Gly-TCC-3 |
| His-GTG | 37 | 1 | His-GTG-2; His-GTG-7 |
| Ile-AAT | 53 | 1 | Ile-AAT-3; Ile-AAT-2; Ile-AAT-17; Ile-AAT-11; Ile-AAT-1; Ile-AAT-12 |
| Ile-TAT | 43 | 1 | Ile-TAT-1; Ile-TAT-5 |
| Leu-CAA | 56 | 1 | Leu-CAA-2; Leu-CAA-8; Leu-CAA-4; Leu-CAA-3; Leu-CAA-15; Leu-CAA-13; Leu-CAA-1; Leu-CAA-16 |
| Leu-CAG | 31 | 1 | Leu-CAG-2; Leu-CAG-4; Leu-CAG-5 |
| Leu-TAA | 66 | 1 | Leu-TAA-3; Leu-TAA-1; Leu-TAA-10; Leu-TAA-8 |
| Leu-TAG/AAG | 65 | 0.58_0.42 | Leu-AAG-1; Leu-AAG-3; Leu-AAG-4; Leu-AAG-6; Leu-TAG-6; Leu-TAG-7; Leu-TAG-4; Leu-TAG-41; Leu-TAG-17; Leu-TAG-2; Leu-TAG-13; Leu-TAG-12 |
| Lys-CTT | 50 | 1 | Lys-CTT-1; Lys-CTT-14; Lys-CTT-2; Lys-CTT-62; Lys-CTT-76 |
| Lys-TTT_Sup-TTA | 35 | 0.89_0.11 | Lys-TTT-6; Lys-TTT-8; Sup-TTA-1 |
| Met-CAT | 61 | 1 | Met-CAT-4; Met-CAT-2; Met-CAT-19; Phe-GAA-1; Phe-GAA-2 |
| Phe-GAA | 45 | 1 | Phe-GAA-1; Phe-GAA-2 |
| Pro-TGG/CGG/AGG | 46 | 0.53_0.32_0.15 | Pro-AGG-1; Pro-CGG-1; Pro-CGG-3; Pro-AGG-7; Pro-TGG-1; Pro-TGG-13; Pro-TGG-3 |
| SeC-TCA | 7 | 1 | SeC-TCA-1 |
| Ser-CGA | 3 | 1 | Ser-CGA-7 |
| Ser-CGA | 17 | 1 | Ser-CGA-14 |
| Ser-GCT | 12 | 1 | Ser-GCT-4 |
| Ser-TGA | 25 | 1 | Ser-TGA-19 |
| Ser-TGA/AGA/CGA | 64 | 0.48_0.3_0.22 | Ser-AGA-9; Ser-AGA-3; Ser-AGA-1; Ser-AGA-2; Ser-CGA-1; Ser-CGA-3; Ser-CGA-5; Ser-CGA-6; Ser-TGA-3; Ser-TGA-5; Ser-TGA-44; Ser-TGA-12; Ser-TGA-1 |

| | | | |
|---|---|---|---|
| Thr-AGT/CGT/TGT | 62 | 0.34_0.33_0.33 | Thr-AGT-3; Thr-AGT-1; Thr-AGT-11; Thr-AGT-28; Thr-AGT-4; Thr-CGT-7; Thr-CGT-6; Thr-NNN-3; Thr-CGT-4; Thr-CGT-1; Thr-CGT-5; Thr-NNN-128; Thr-AGT-7; Thr-NNN-90; Thr-TGT-1; Thr-TGT-4 |
| Thr-CGT | 8 | 1 | Thr-NNN-15 |
| Thr-TGT | 51 | 1 | Thr-TGT-7; Thr-TGT-8 |
| Trp-CCA | 11 | 1 | Trp-CCA-4 |
| Trp-CCA | 57 | 1 | Trp-CCA-3; Trp-CCA-8; Trp-CCA-1; Trp-CCA-2 |
| Tyr-GTA | 39 | 1 | Tyr-GTA-1; Tyr-GTA-2 |
| Val-AAC | 41 | 1 | Val-AAC-30; Val-AAC-6 |
| Val-CAC/TAC/AAC | 55 | 0.77_0.14_0.09 | Val-AAC-2; Val-AAC-1; Val-CAC-1; Val-CAC-11; Val-CAC-2; Val-CAC-4; Val-CAC-6; Val-CAC-8; Val-TAC-2; Val-TAC-1 |
| Val-TAC | 13 | 1 | Val-TAC-22 |
| Val-TAC | 38 | 1 | Val-TAC-4; Val-TAC-10 |
| iMet-CAT | 36 | 1 | iMet-CAT-1; iMet-CAT-11; iMet-CAT-2 |
| mt-Ala-TGC | 27 | 1 | mt-Ala-TGC |
| mt-Arg-TCG | 30 | 1 | mt-Arg-TCG; mt-Arg-TCGs |
| mt-Asn-GTT | 30 | 1 | mt-Asn-GTT |
| mt-Asp-GTC | 23 | 1 | mt-Asp-GTC; mt-Asp-GTCs |
| mt-Cys-GCA | 29 | 1 | mt-Cys-GCA |
| mt-Gln-TTG | 29 | 1 | mt-Gln-TTG |
| mt-Glu-TTC | 1 | 1 | mt-Glu-TTC |
| mt-Gly-TCC | 24 | 1 | mt-Gly-TCC |
| mt-His-GTG | 0 | 1 | mt-His-GTG |
| mt-Ile-GAT | 19 | 1 | mt-Ile-GAT |
| mt-Leu1-TAG | 9 | 1 | mt-Leu1-TAG |
| mt-Leu2-TAA | 21 | 1 | mt-Leu2-TAA |
| mt-Lys-TTT | 2 | 1 | mt-Lys-TTT |
| mt-Met-CAT | 15 | 1 | mt-Met-CAT |
| mt-Phe-GAA | 14 | 1 | mt-Phe-GAA |
| mt-Pro-TGG | 10 | 1 | mt-Pro-TGG |
| mt-Ser1-GCT | 18 | 1 | mt-Ser1-GCT |
| mt-Ser2-TGA | 20 | 1 | mt-Ser2-TGA; mt-Ser2-TGAs |
| mt-Thr-TGT | 4 | 1 | mt-Thr-TGT; mt-Thr-TGTs |
| mt-Trp-TCA | 34 | 1 | mt-Trp-TCA |
| mt-Tyr-GTA | 34 | 1 | mt-Tyr-GTA |
| mt-Val-TAC | 48 | 1 | mt-Val-TAC |

**Supplementary Table 2 Detected modifications per cluster.**

Modifications inferred from RT-signature in mock samples and C retention in BS samples.

| | $m^1G9$ | $m^1A9$ | $m^1A14$ | $acp^3U$ 20/20a/20 | $m^3C20$ | $m^{2,2}G$ 26/27 | $m^3C32$ | I34 | $m^5C34$ | $m^1G37$ | $m^1I37$ | $o^2yW37$ | $ms^2t^6A37$ | $ms^2i^6A37$ | $m^5C38$ | $m^5C40$ | $m^3Ce2$ | $m^5C$ 48/49/50 | $m^1A58$ | $m^5C72$ |
|---|---|---|---|---|---|---|---|---|---|---|---|---|---|---|---|---|---|---|---|---|
| Ala-AGC | | | | | | + | | + | | | + | | | | | | | | + | |
| Ala-TGC/CGC | | | | + | | + | | | | | + | | | | | | | + | + | |
| Arg-ACG | + | | | | | + | | + | | + | | | | | | | | + | + | |
| Arg-CCT | + | | | | | | + | | | | | | | | | | | | + | |
| Arg-TCG | + | | | | | + | | | | + | | | | | | | | | + | |
| Arg-TCG | + | | | | | + | | | | + | | | | | | | | | + | |
| Arg-TCG/CCG | + | | | | | | | | | + | | | | | | | | | + | |
| Arg-TCT | + | | | | | | + | | | | | | | | | | | | + | |
| Asn-GTT | + | | | | | + | | | | | | | | | | | | | + | |
| Asp-GTC | | + | | | | | | | | | | | | | + | | | +/+ | | |
| Cys-GCA | | | | + | | | | | | + | | | | | | | | + | + | + |
| Gln-CTG/TTG | + | | | | | | | | | | | | | | | | | +/+ | + | |
| Glu-CTC | | | | | | | | | | | | | | | | | | +/+ | + | |
| Glu-TTC/CTC | + | | | | | | | | | | | | | | | | | +/+ | + | |
| Gly-CCC | | | | | | | | | | | | | | | | | | +/+ | + | |
| Gly-GCC/CCC | | | | | | | | | | | | | | | + | + | | +/+/+ | + | |
| Gly-TCC | + | | | | | | | | | | | | | | | | | +/+/+ | + | |
| His-GTG | | | | | | | | | | + | | | | | | | | + | + | |
| Ile-AAT | | | | + | | + | | + | | | | | | | | | | + | + | |
| Ile-TAT | + | | | | | + | | | | | | | | | | | | | + | |
| Leu-CAA | | | | + | | | | | + | + | | | | | | | | + | + | |
| Leu-CAG | | | | + | | + | | | | + | | | | | | | + | + | + | |
| Leu-TAA | | | | | | + | | | | + | | | | | | | | + | + | |
| Leu-TAG/AAG | | | | + | | + | | + | | + | | | | | | | | + | + | |
| Lys-CTT | | | | | | | | | | | | | | | | | | + | + | |
| Lys-TTT_Sup-TTA | | | | | | | | | | | | | + | | | | | +/+ | + | |
| Met-CAT | + | | | + | | + | | | | | | | | | | | | + | + | |
| Phe-GAA | | | + | | | + | | | | | | + | | | | | | +/+ | + | |
| Pro-TGG/CGG/AGG | + | | | | | | | + | | + | | | | | | | | +/+ | + | |
| SeC-TCA | | | | | | | | | | | | | | | | | | | + | |
| Ser-CGA | | | | | | + | + | | | | | | | | | | | + | + | |
| Ser-CGA | | | | | | + | + | | | | | | | | | | | + | + | |
| Ser-GCT | | | | | | + | + | | | | | | | | | | + | + | + | |
| Ser-TGA | | | | | | + | + | | | | | | | | | | + | + | + | |
| Ser-TGA/AGA/CGA | | | | | | + | + | + | | | | | | | | | + | + | + | |
| Thr-AGT/CGT/TGT | + | | | | | + | + | + | | | | | | | | | | +/+ | + | + |
| Thr-CGT | + | | | + | | + | + | | | | | | | | | | | + | + | + |
| Thr-TGT | | | | + | | | + | | | | | | | | | | | + | + | + |
| Trp-CCA | + | | | | | + | | | | + | | | | | | | | | + | |
| Trp-CCA | + | | | | | + | | | | + | | | | | | | | | + | |
| Tyr-GTA | | | | + | | +/+ | | | | + | | | | | | | | | + | |
| Val-AAC | | | | | | + | | + | | | | | | | | | | +/+ | + | |
| Val-CAC/TAC/AAC | | | | | | | | + | | | | | | | | | | +/+ | + | |
| Val-TAC | | | | | | | | | | | | | | | | | | +/+ | + | |
| Val-TAC | | | | | | | | | | | | | | | | | | +/+ | + | |
| iMet-CAT | + | | | | | + | | | | | | | | | | | | + | + | |
| mt-Ala-TGC | | + | | | | | | | | | | | | | | | | | | |

| | | | | | | | | | | | | | | | | | | | |
|---|---|---|---|---|---|---|---|---|---|---|---|---|---|---|---|---|---|---|---|
| mt-Arg-TCG | | + | | | | | | | | + | | | | | | | | | |
| mt-Asn-GTT | | | | | | + | | | | | | | | | | | | | + |
| mt-Asp-GTC | | + | | | | | | | | | | | | | | | | | + |
| mt-Cys-GCA | + | | | | | | | | | | | | | | | | | | + |
| mt-Gln-TTG | + | | | | | | | | | | | | | | | | | | + |
| mt-Glu-TTC | | + | | | | | | | | + | | | | | | | | | + |
| mt-Gly-TCC | | + | | | | | | | | | | | | | | | | | |
| mt-His-GTG | | + | | | | | | | | | | | | | | | | | + |
| mt-Ile-GAT | + | | | | | + | | | | + | | | | | | | | | + |
| mt-Leu1-TAG | | + | | | | | | | | + | | | | | | | | + | + |
| mt-Leu2-TAA | + | | | | | | | | | + | | | | | | | | + | + |
| mt-Lys-TTT | | | | | | + | | | | | | | | | | | | | |
| mt-Met-CAT | | | | | | | | | + | | | | | | | | | | + |
| mt-Phe-GAA | | + | | | | | | | | + | | | | | | | | | |
| mt-Pro-TGG | | + | | | | | | | | + | | | | | | | | | + |
| mt-Ser1-GCT | | | | | | | | | | | | | | | | | | | + |
| mt-Ser2-TGA | + | | | | | + | + | | | | | | | + | | | | | + |
| mt-Thr-TGT | | + | | | | | + | | | | | | | | | | | + | + |
| mt-Trp-TCA | | + | | | | | | | | | | | | + | | | | | + |
| mt-Tyr-GTA | + | | | | | + | | | | + | | | | | | | | + | + |
| mt-Val-TAC | + | | | | | | | | | | | | | | | | | | + |

**Supplementary Table 3  Reads pre-processing summary.**

Number of reads that successfully passed a given preprocessing step

| | raw reads | adapter trimmed reads | size selection (>=29nts) | remaining after 5’ polyT trimming + size selection (>=20nts) | fastp qc passed reads |
|---|---|---|---|---|---|
| mean | 4902502 | 4668533 | 3499983 | 3485431 | 3473257 |
| min | 1723083 | 1595365 | 507173 | 498841 | 496495 |
| max | 18460912 | 17098130 | 11960487 | 11820385 | 11737789 |
| fraction remaining | 1 | 0.952 | 0.714 | 0.711 | 0.708 |

### SUPPLEMENTARY FIGURES

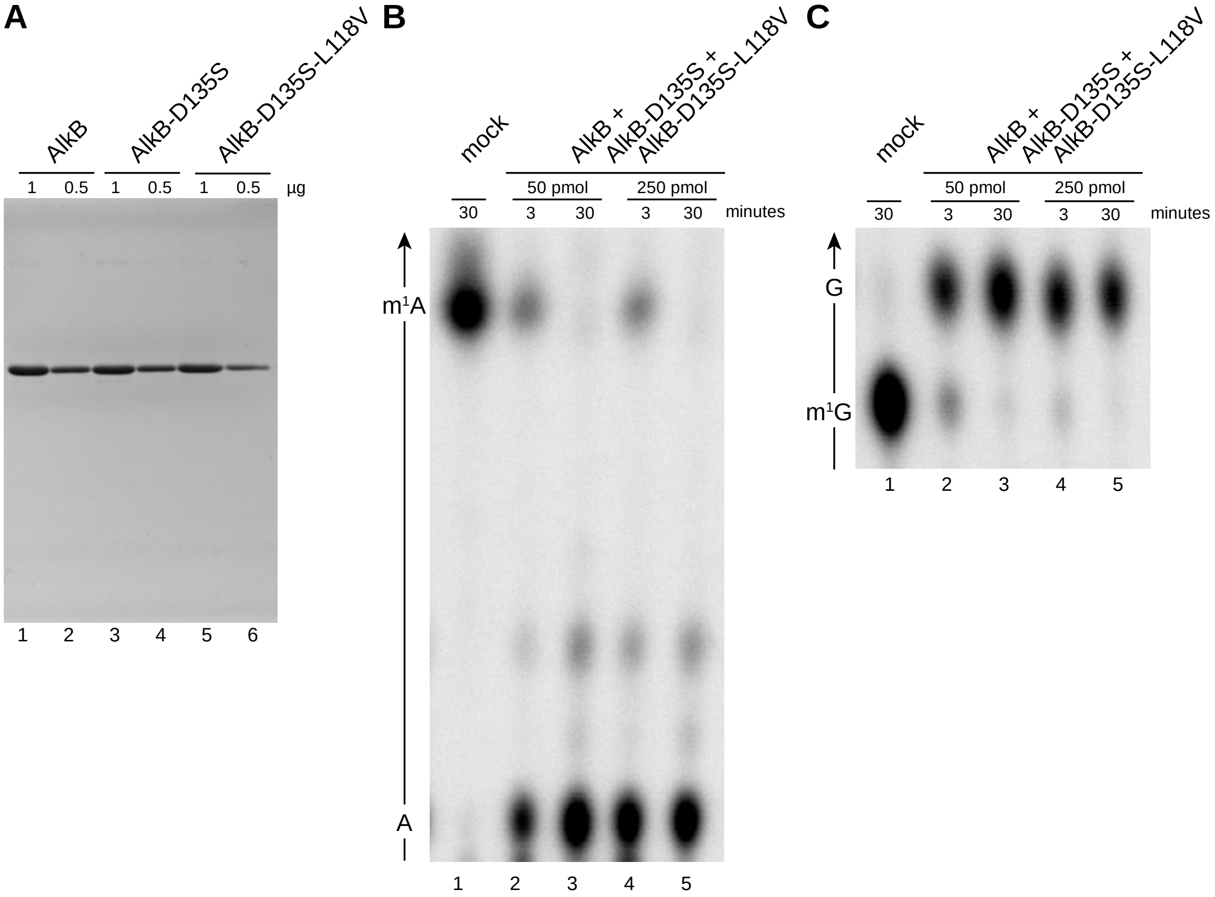

**Supplementary Figure 1 AlkB purification and activity test**. (**A**) Purity assessment of AlkB (and its mutants) by SDS-PAGE. On top the AlkB form and the amount loaded is annotated. (**B**) and (**C**) AlkB demethylation assay on RNA oligonucleotides carrying a 5' $^{32}$P-labelled m$^1$A (B) or m$^1$G (C). The oligoes were incubated at varying concentrations of AlkB mix and reaction times. After P1 digestion, the nucleoside monophosphate residues were resolved by TLC. Arrows on the left indicate the direction of the solvent migration and highlight the nucleotide identity.

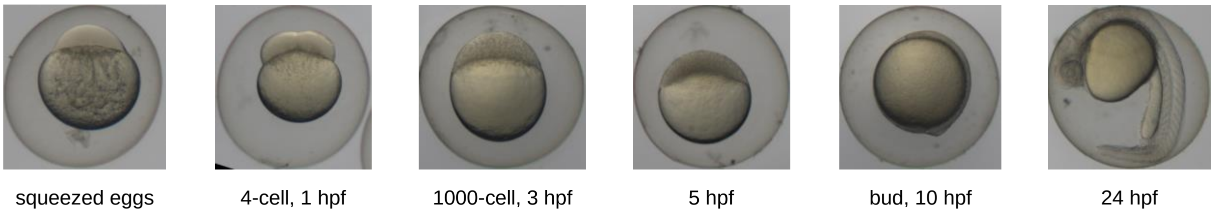

**Supplementary Figure 2 Zebrafish embryo development stages.** Representative microscopy images of the unfertilised eggs and embryo developmental stages used in the study. Labels indicate the stage description, and time in terms of hours post fertilization (hpf).

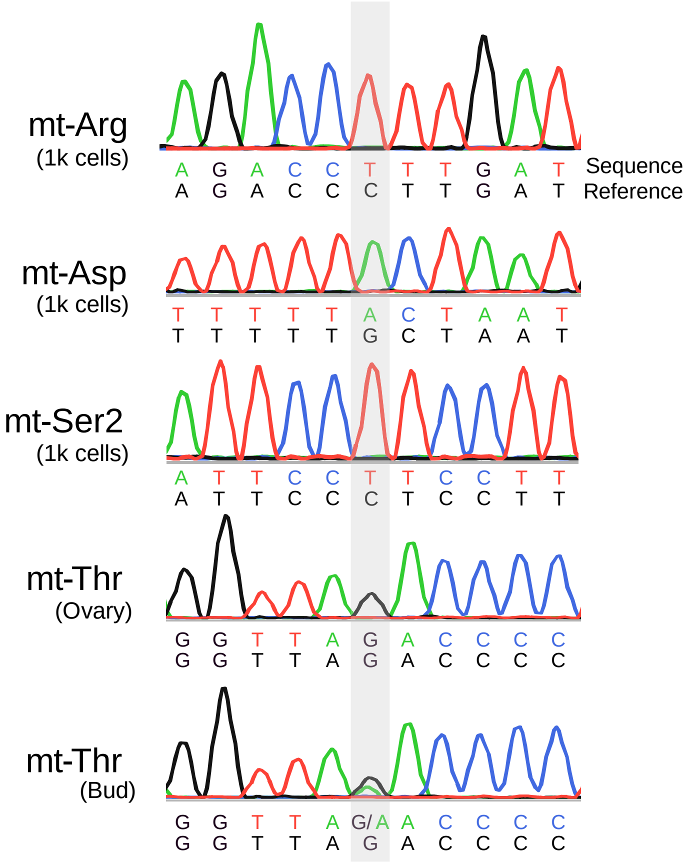

**Supplementary Figure 3 Sanger sequencing analysis of mitochondrial DNA.** Sanger sequencing results of amplified mitochondrial DNA from the samples indicated are shown as chromatogram traces. The obtained sequences were aligned to the reference zebrafish mitochondrial tRNA sequence from mitotRNAdb. SNPs identified are emphasized with a grey background.

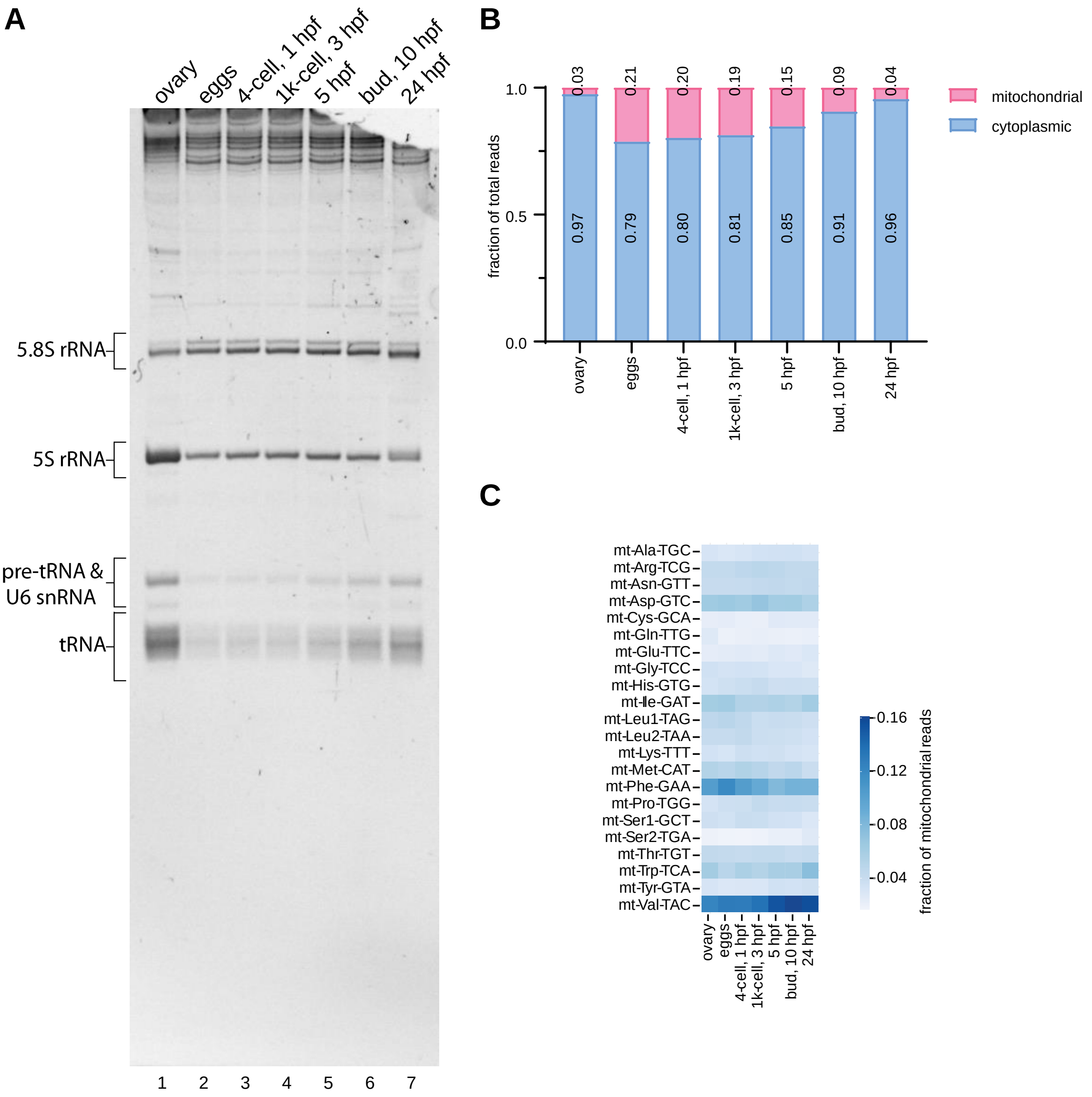

**Supplementary Figure 4 Small RNA expression during zebrafish embryo development**. (**A**) Total RNA extracted from zebrafish ovary, unfertilised eggs, and embryos at different developmental stages was separated on a denaturing 15% polyacrylamide gel. The sample identity is indicated on the top; on the left the main small RNA bands are identified. (**B**) Relative abundance of mitochondrial versus nucleo-cytoplasmic tRNAs, expressed as fraction of total reads. Data are means of three biological replicates with the exception of the activated eggs (n=2). (**C**) Heat-map representing the normalized abundance of the mitochondrial tRNAs (y-axis) in the analysed samples (x-axis). The number of reads mapped to each tRNA was normalized by the total mitochondrial tRNAs read count and plotted. The blue colour scale indicates the fraction of reads mapped to the individual tRNA. Data are means of three biological replicates with the exception of the activated eggs (n=2).

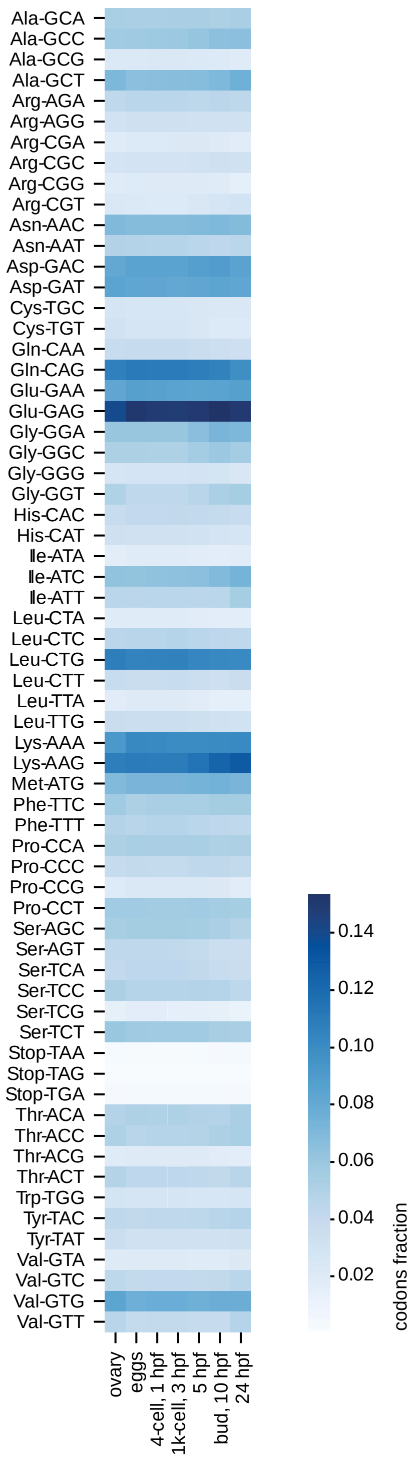

**Supplementary Figure 5 Codon frequency dynamics during zebrafish embryo development**. Heat-map showing the codon frequency in the transcriptome across zebrafish ovary, unfertilised eggs, and embryo developmental stages (x-axis). Codons and amino acids are indicated on the y-axis. The blue colour scale indicates the codon fraction calculated as follows: for each protein coding gene, the occurrences of each codon was counted and multiplied by the genes expression level (normalized read count). To obtain the codon frequency, those per codon counts were summed up over all genes and then divided by the total number of codons.

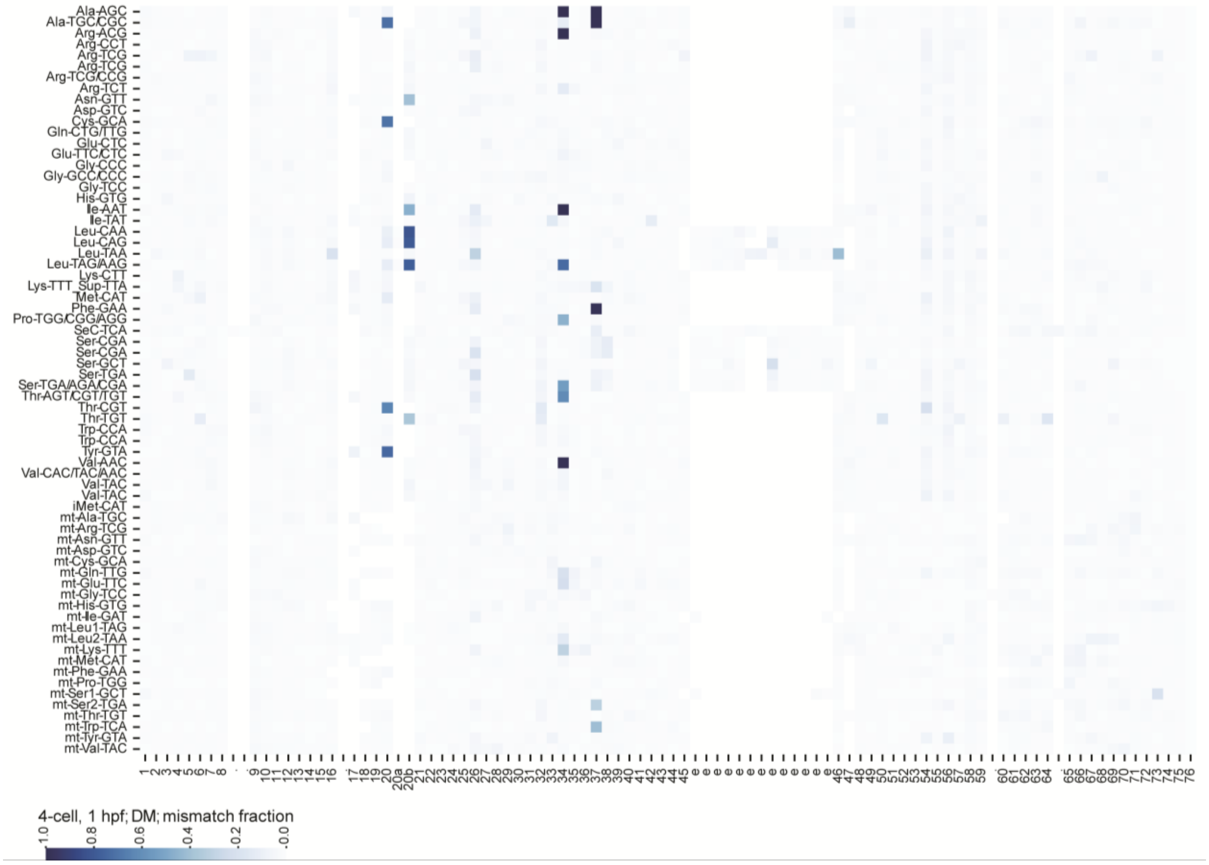

**Supplementary Figure 6 Misincorporation signatures of demethylated tRNA from 4-cell stage zebrafish embryos**. Heat-map of misincorporation fraction of all cytosolic tRNA clusters and mt-tRNAs (y-axis). Canonical nucleotide positions are annotated on the x-axis. Nucleotide positions that are rarely present in the tRNA clusters are annotated with a dot. The blue colour scale indicates the mean mismatch fraction across three biological replicates.

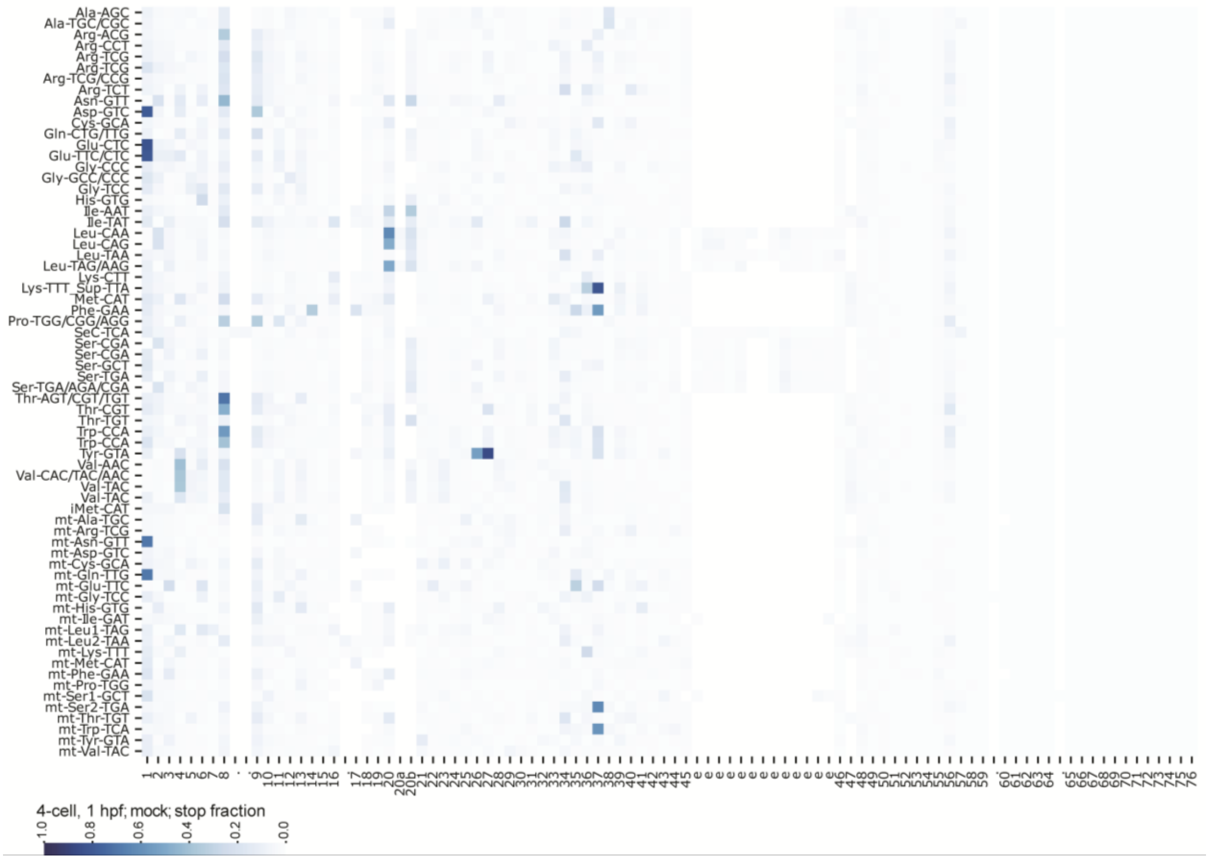

**Supplementary Figure 7 RT stop signatures of tRNA from 4-cell stage zebrafish embryos**. Heat-map of RT stop fraction of all cytosolic tRNA clusters and mt-tRNAs (y-axis). Canonical nucleotide positions are annotated on the x-axis. Nucleotide positions that are rarely present in the tRNA clusters are annotated with a dot. The blue colour scale indicates the mean RT stop fraction across three biological replicates. The stop signal at the 5' end of the clusters Asp-GTC, Glu-CTC, Glu-TTC/CTC, mt-Asn-GTT and mt-Gln-TTG are most likely caused by over-trimming of leading Ts (see experimental details in Supplementary File 1).

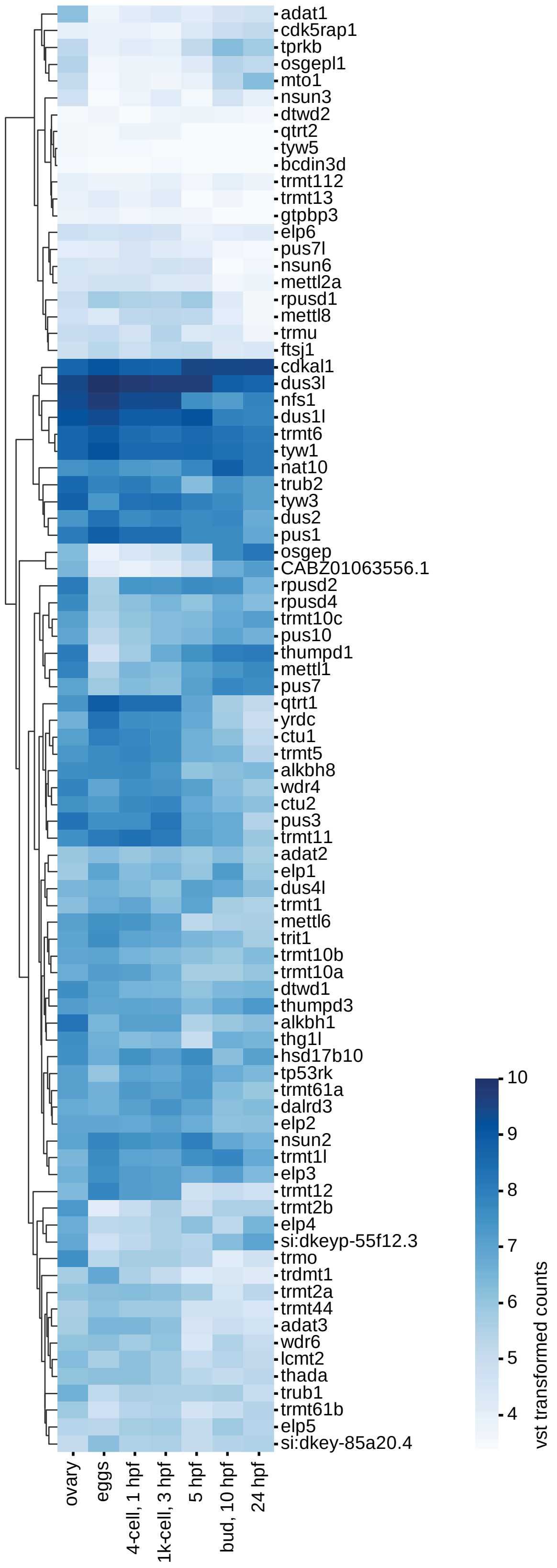

**Supplementary Figure 8 Gene expression analysis of tRNA modification enzymes**. Heat-map showing mean VST transformed counts (blue colour scale) for tRNA modification enzymes (y-axis) across zebrafish ovary and embryo development (x-axis). The tRNA modification enzymes were hierarchically clustered based on their similarity in expression levels or based on their similarity in expression dynamics.

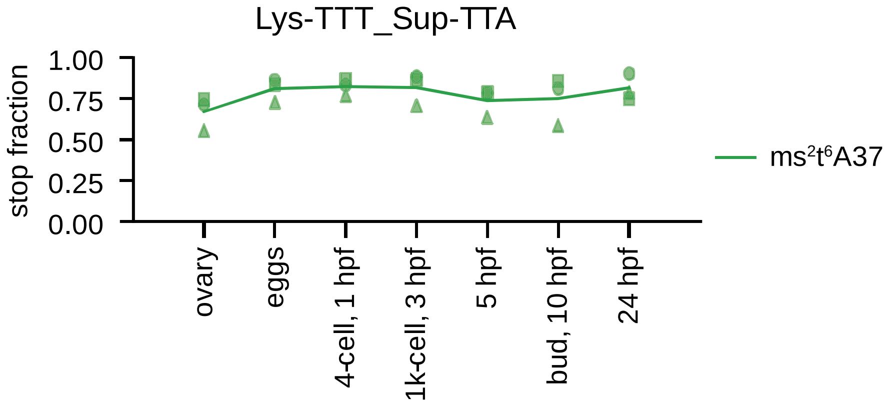

**Supplementary Figure 9 RT stop caused by ms$^2$t$^6$A37 in the cluster Lys-TTT_Sup-TTA.** RT stop fraction (y-axis) at position A37 in the cluster Lys-TTT_Sup-TTA across zebrafish embryo development (x-axis). The line represents the mean of three biological replicates, which are indicated by square, circle and triangle.

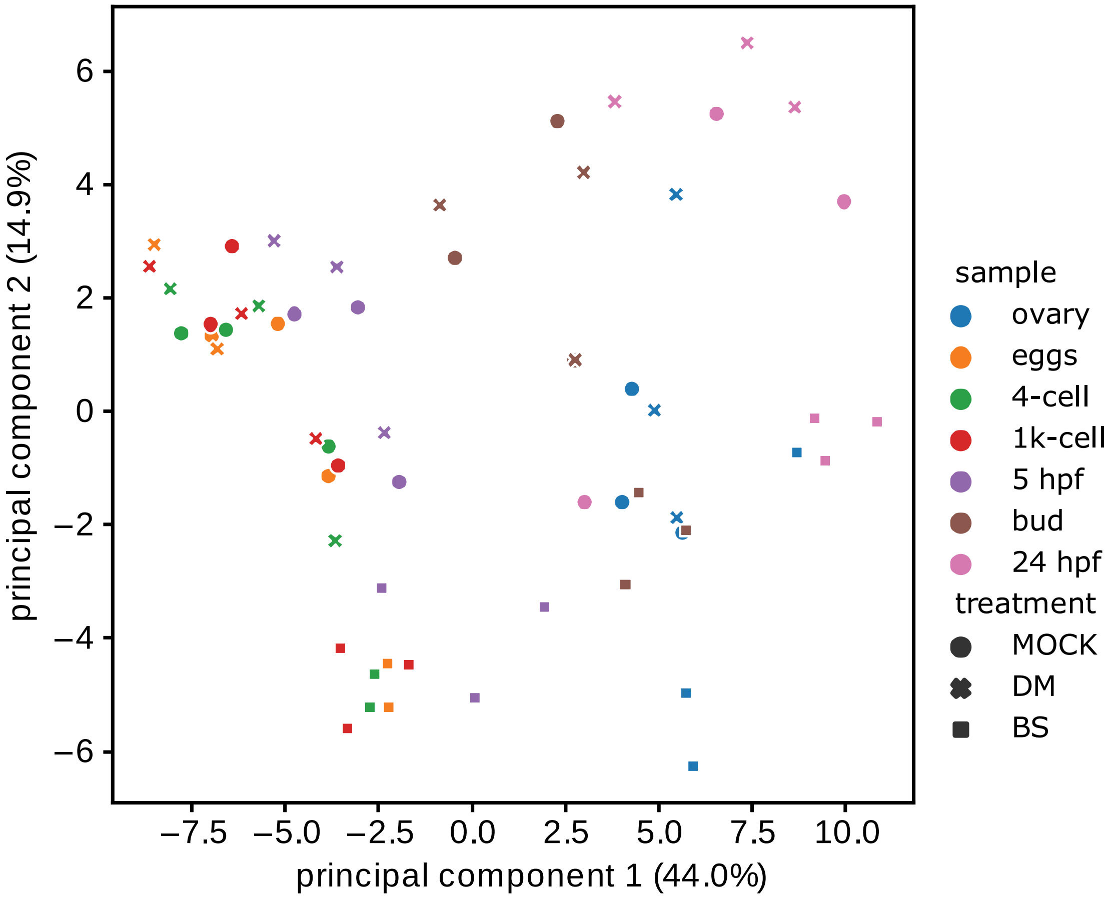

**Supplementary Figure 10 PCA analysis of tRNA sequencing data.** Plot of the first two dimensions of a principal component analysis, based on the normalized abundance (in RPM) of each tRNA cluster. Samples and treatments are identified by colors and symbols according to the legend.

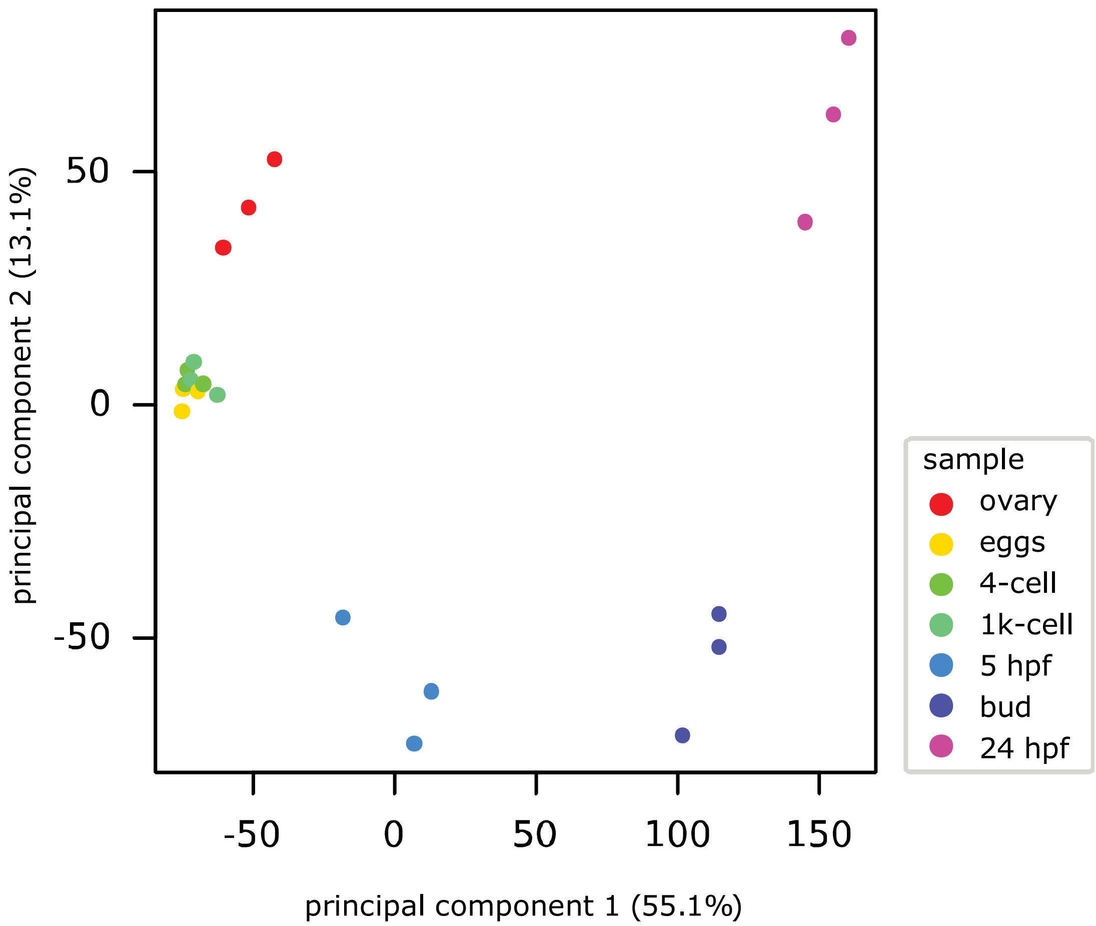

**Supplementary Figure 11 PCA analysis of mRNA sequencing data**. Plot of the first two dimensions of a principal component analysis, based on the transformed count data. Samples are identified by colors according to the legend.
